# Methionine metabolism and the NOP2 methyltransferase are essential for MYC-Driven liver tumorigenesis

**DOI:** 10.64898/2026.01.28.702329

**Authors:** Sensen Lin, Charles Berdan, Moriah Sandy, Xinjun Lu, Vijay Ramani, Daniel K. Nomura, Xin Chen, Joyce Lee, Andrei Goga

## Abstract

Hepatocellular carcinoma (HCC) represents the third leading cause of cancer-related death worldwide and has been increasing in developed nations.^1,2^ The MYC oncogene or its paralogs are frequently amplified or overexpressed in subtypes of cancer associated with stem cell-like features and worse clinical outcomes,^3,4^ including in liver cancer.^5^ Unfortunately, selective inhibitors that target MYC or its transcriptional program are not yet clinically available for therapy of HCC. Here, we identified methionine metabolism as a selective vulnerability for MYC but not RAS-driven liver cancers. MYC-driven liver cancer cells are methionine dependent, with markedly diminished tumor growth when mice are fed a methionine low diet. While RAS-driven liver cancer was resistant to a low methionine diet. S-adenosylmethionine (SAM), the predominant methyl donor, partially rescues cell proliferation following methionine depletion, suggesting that methylation processes are especially critical in the context of MYC high tumor cells. Heavy isotope methionine tracing in MYC high cells identified increased levels of m5C nucleotides. We found NOP2, an rRNA m5C-methyltransferase, was regulated by both MYC overexpression and methionine abundance linking the two processes. Methionine depletion reduced methylation of multiple 28S rRNA residues as did NOP2 knockdown. Depletion of NOP2 selectively inhibited MYC liver cancer cell proliferation and *in vivo* tumor growth. Thus, methionine catabolism is critical for MYC-driven liver tumorigenesis and the rRNA methyltransferase NOP2 may serve as a new therapeutic target in liver cancer.

## Introduction

MYC, a basic helix-loop-helix zipper (bHLHZ) motif– containing transcription factor, is one of the most commonly amplified genes in human cancer.^6^ MYC can activate or repress hundreds of target genes,^7,8^ including genes transcribed by all three major polymerases. Likewise, MYC overexpression can broadly alter genome organization and epigenetics by cooperating with other transcription factors and epigenetics regulators; it alters chromatin 3D configuration, changes topologically associating domains, and regulates gene expression at long-range.^9,10^ MYC also induces alterations of key processes such as multiple cellular metabolic processes, protein synthesis, RNA metabolism, and mitochondrial and ribosome biogenesis,^11-14^ resulting in increased cell proliferation, protein synthesis, DNA replication, and global reprogramming of cellular metabolism. We postulated that as MYC reprograms liver cancer metabolism,^15,16^ it can render tumors dependent on specific nutrients.

Methionine is one of nine essential amino acids required for growth and development.^17,18^ Humans obtain methionine from food and gastrointestinal microbes. Methionine is unique among amino acids, as it contains a sulfur and methyl group, which are essential for biological processes such as methylation, anti-oxidation and maintenance of protein structure.^19,20^ Cancer cells often have disrupted methionine metabolism and altered methylation.^21,22^ Several types of cancers have been found to have a dependence on methionine,^23,24^ while non-tumor normal cells can be grown in the absence of methionine but with supplementation by the precursor homocysteine.^25^ Methionine restriction along with, or in combination with, chemotherapeutics, shows anti-cancer effects against chemo-resistant or radiation-resistant cancers.^26,27^ However, which oncogene-driven tumors require methionine, what is the mechanism of methionine dependence, and which pathways downstream of methionine metabolism could be targeted in cancer remains poorly understood. In this study, we examine alterations in the methionine pathway in MYC-driven liver cancer, identifying a unique dependency on NOP2, an RNA methyltransferase with a role in ribosome biogenesis. This finding has implications for the development of future cancer therapeutics for this difficult to treat cancer type.

## Methods

### LT2-MYC transgenic model

The LT2-MYC transgenic mice were previous described.^28^ MYC transgene segregates with Y chromosome, such that only male mice were used for the research. At 6 weeks of age, mice were fed with doxycycline free diet and carefully monitored for liver tumor development.

All animal experiments were performed in compliance with approved protocols by the Institutional Animal Care and Use Program at UCSF. All mice were housed under a 12 h light -12 h dark cycle. Room temperature mice were housed at 23℃in ventilated cages. Unless otherwise stated, mice were fed a standard diet (Lab Diet 5008) and had free access to food and water.

### Cell culture

Murine EC4 cells were described previously.^29^ Cells were maintained in high glucose DMEM supplemented with 10% FBS and 1×non-essential amino acid and sodium pyruvate. To turn off MYC expression, 10 ng/ml doxycycline was added. Human HCC cell lines Hep40, SNU398, PLC/PRF/5 and SNU475 were a gift from Dr. Xin Chen at University of California, San Francisco. HepG2 cells was purchased from ATCC. Hep40, SNU398, PLC/PRF/5 and HepG2 cells were cultured in DMEM supplemented with 10% FBS. SNU475 cells were maintained in RPMI-1640 added with 10% FBS. All the cell lines were tested for mycoplasma contamination before experiments. For methionine restriction assay, cells were suspended in methionine free medium and methionine supplementation to indicated concentrations were added.

### Cell proliferation assay

Cell proliferation was assessed by automatically counting cell numbers using Biotek Cytation 5 high content microscope according to the manufacturer’s instructions. Briefly, cells were seeded at 1000 cells per well in 96 well plates. After treated for the indicated time, cells were washed, fixed with 3.7% formaldehyde and stained with 5µg/ml Hoechst 33342 for 30 minutes. After washing, the plates were placed in the microscope, images were captured under the DAPI filter cube, and cell numbers were determined by Gen5 automated counting on the images.

### Cell cycle analysis

EC4 cells were treated with methionine free media or normal DMEM (200µM methionine) for 1-5 days, after that cells were harvested and fixed by 70% ethanol overnight. The next morning, cells were washed with PBS for 3 times, and stained with 50 µg/ml propidium iodide solution at room temperature for 30 minutes. 10^5^ cells were analyzed with BD flow cytometry.

### Metabolite rescue assay

EC4 cells were plated at 500 cells per well in 96 well plates; the next morning cells were washed 3 times with PBS and DMEM media containing 1µM methionine were added. Methionine pathway-related metabolites were added back at a concentration of 100µM. Cells were cultured for 1-7 days and cell number was counted as described above.

### Western blot

Total cellular extracts were prepared using standard procedures. Cells were lysed with RIPA supplemented with protease inhibitor cocktail and phosSTOP. Protein concentrations were measured with Pierce BCA Protein Assay Kit. The primary antibodies used in this paper were: β-actin (1:10000), c-MYC (1:1000), c-MYC S62 phosphorylation (1:2000), MTR (1:900), MAT1A (1:1000), MAT2A (1:2000), NOP2 (1:1000), Vinculin (1:1000), MTAP (1:1000), CBS (1:1000), SHMT2 (1:1000).

### RNA interference

The RNAi method was described in our previous work.^30^ ON-TARGETplus siRNAs (a mixture of 4 siRNAs) for human MYC (L-003282-02-005), mouse Nop2 (L-053142-01-0005), human NOP2 (L-020084-00-0005) as well as Non-target control (D-001810-10-20) were purchased from Dharmacon (SMARTpool). Briefly, cells were treated with 10-30nM specific siRNAs for 24-72 h, using Lipofectamine RNAiMAX transfection regent following the manufacturer’s manual. Knockdown effect was confirmed by western blotting.

### Lentivirus production and infection

HEK 293T cells were seeded in 6 well plates, 6×10^5^ per well. The next morning, cell medium was changed with fresh DMEM without antibiotics. Add 6µl polyethylenimine (PEI, 1 mg/ml) to 100µl pre-warmed OPTI-MEM; add 2µg total DNA to 100µl pre-warmed OPTI-MEM (psPAX2:pMD2.G:pDECKO=3.5:1.5:5). Add diluted PEI to diluted DNA, and incubate the mixture for 30 min at room temperature. Gently transfer the mixture to 293T cells, and incubate the cells for another 48 h.

48 h after transfection, collect lentivirus from cell culture supernatants. Spin the supernatants at 2000rpm for 10 min to get rid of cell debris. Syringe filter (0.45µm) the supernatants, and transduce EC4 cells with fresh virus.

Apply 200µl fresh virus to 3×10^10^ EC4 cells in 10cm cell culture dish. 48 h after infection, change cell medium and add puromycin (2µg/ml) to select for infected cells. 3 days after selection, harvest cells for western blot analysis.

### Metabolic analysis

Metabolic analysis was performed as previously described.^29^ Samples were extracted according to Metabolon’s standard solvent extraction method. LC-MS/MS was performed and d3N15 serine was used as the internal control.

### Methionine tracing

EC4 cells were plated in 6 well plates, the next day cell culture medium was changed to contain L-methionine- (methyl-13C), either high methionine (200 µM) or low methionine (10 µM). Cells were cultured for 6-12h. After that culture media was aspired, and cells were washed with ice cold ammonium acetate wash solution twice. The wash solution was discarded; cells were incubated with an 80% methanol solution. Plates were placed in -80°C freezer for 15 min. Following freezing, adherent cells were scraped off and cell solutions were centrifuged at 17,000g for 10 min at 4°C. The top 250 µl supernatant was transferred to a new tube and dried under nitrogen and vacuum (Turbovac). Dried samples were reconstituted in 70 µl LCMS grade water to dissolve the samples for LC-MS analysis.

Untargeted LCHRMS analysis of cell extracts was performed on a Sciex Exion UPLC equipped with a Kinetex F5 column (Phenomenex, 150 x 2.1 mm, 2.6 um) and coupled to a Sciex TripleTOF 6600+ operated in negative mode electrospray ionization with IDA acquisition of MS2 (accumulation time 0.25 s, collision energy 30V (15V spread), acquisition window 50-1000Da). A gradient of 0% - 95% acetonitrile over 12 min was used for compound separation.

Raw data from the TripleTof mass spectrometer were converted to mzXML format for further analysis. Peak finding, filtering, alignment, scaling, and identification were performed using open-source Scripps XCMS Online platform (https://xcmsonline.scripps.edu/). Then, the data matrix comprising sample identity, ion identity (retention time, RT, and m/z) and relative ion abundance were imported into SIMCA-P+TM software^31^ for multivariate data analysis. Supervised projection to latent structures-discriminant analysis (PLS-DA) and orthogonal partial least-squares discriminant analysis (OPLS-DA) were performed to identify metabolites increased with labeled-methionine medium culture time. Among these metabolites, isotope-labeled metabolites were further pinpointed by picking metabolites having both unlabeled and labeled isotopologues, with labeled isotopologues showing cultured time-dependent increase.

### Chromatin Immunoprecipitation (ChIP)

3×10^10^ EC4 cells were plated in 15cm cell culture dish, 48 h later culture medium was discarded and add 1% paraformaldehyde to fixate the cells for 10 min. The fixation was quenched by 0.125 M glycine, and cells were washed with PBS twice. 5ml cell lysis buffer (10mM Tris-HCl (pH8.1), 10mM NaCl, 1.5mM MgCl_2_, 0.5% Igepal-CA630) was added and cells were scraped into 15ml conical tubes. Cells were lysed by rotation in 4°C for 15 min. After centrifuging, samples were resuspended in 0.5ml nuclear lysis buffer (50mM Tris-HCl, 5mM EDTA, 1% SDS). Samples were then sonicated at 4°C for 30 min. Insoluble components were removed by spin at 13000 RCF for 15 min at 4°C. Supernatants were collected and mixed with 0.5ml of dilution buffer (16.7mM Tris-HCl, 1.1% Triton X-100, 0.01% SDS, 167mM NaCl, 1.2mM EDTA). Lowry method was employed to quantify total protein. 500 µg total protein was used for IP. 2µg MYC antibody (Abcam, ab32072) and 20µl Millipore protein A/G beads were used for each IP.

The next day, remove the supernatants by putting samples on a magnetic rack. Samples were washed in the following order: low salt buffer (0.1% SDS, 1% Triton X-100, 2mM EDTA, 20mM Tris-HCl, 150mM NaCl), high salt buffer (0.1% SDS, 1% Triton X-100, 2mM EDTA, 20mM Tris-HCl, 500mM NaCl), LiCl buffer (1% Igepal-CA630, 1% deoxycholate, 1mM EDTA, 10mM Tris-HCl, 250mM LiCl), TE buffer (10mM Tris-HCl, 1mM EDTA).

After removal of TE buffer, elution buffer (10mM Tris-HCl, 1mM EDTA, 1% SDS, 150mM NaCl, 5mM DTT) was added and samples were incubated at 65°C for 10 min. Collect the elute. Samples were incubated at 65°C overnight to reverse crosslinking.

Proteinase K (20mg/ml) was used to digest samples. Active Motif DNA purification kit (58002) was used to purify DNA. SYBR Green Master Mix kit (A25742, Applied Biosystems) was used to detect MYC binding sites on target genes. Quantitative PCR primers for genomic regions for *Nop2*: forward 5’-CGGTGTGCGACCTCTGAGTTC-3’, reverse 5’-TCCTGCAGACGCACCTGAAG-3’; *Odc1* forward 5’-ATCCATCCTCCGCTTGCATCT-3’, reverse 5’-CTTCCTCGGCTTTCCATTCAG-3’; Negative forward 5’-TGTCCCATTCACGTGTTAGATG-3’, reverse 5’-GGAACCCTTGGTCTCCTCAT-3’.

### Click-iT HPG kit Alexa Fluor 488 Protein Synthesis Assay

Protein synthesis assay was performed following the manufacturer’s manual. Briefly, 1000 cells were seeded into 96 well plates, 48 h later cells were fixated with 3.7% formaldehyde, followed by permeabilization with 0.5% Triton® X-100. After PBS washing, cells were incubated with 100 µl/well Click-iT reaction solutions for 30 min. Cell nucleus was stained with 5µg/ml Hoechst for another 30 min, and then photographed with Biotek Cytation 5. Cell fluorescence intensity was analyzed with Gen5 software.

### Generation of Nop2 KD cell lines

*Nop2* KO/KD cell lines were generated with CRISPR technique. EC4 cells were transduced with lentiCas9 virus and selected with 10 µg/ml blasticidin for 7 days; Cas9 expression was confirmed by western bolt. *Nop2* sgRNA sequences: 5’-TGCATAGATGTGAGCTGTTA-3’ and 5’- CAACACAGTGATGTCACCAA-3’ were cloned into pDECKO-mCherry plasmid (Addgene #78534) according to previous report.^32^ The pDECKO-*Nop2* sgRNAs plasmid was validated with Sanger seq. EC4-Cas9 cells were infected with pDECKO-*Nop2* sgRNAs virus. 48 h after transfection, cells were selected with puromycin for another 3 days. Cells were then sorted into 96 well plates with mCherry, and maintained until single cell clones appeared. *Nop2* DNA mutation was confirmed by western blot and sanger seq.

### RNA-seq

MYC^high^ and MYC^low^ EC4 cells were seeded at 6 well plates, 3×10^10^ per well. The next morning, cell culture media was replaced and cells were washed with PBS and treated with either low methionine DMEM (10µM) or high methionine media (200µM) for 24 h. After treatment, cells were harvested and total RNA was isolated according to the manufacturer’s protocol (RNeasy Plus Mini kit, Qiagen). RNA quality was measured on Agilent 2100 Bioanalyzer. Samples were sent to Novogen for library construction and sequencing. Reads quality was viewed by FastQC, and raw data were then trimmed by Fastp with parameters -- detect_adapter_for_pe -q 20 -u 50 -n 5. Trimmed reads were then mapped to mouse genome GRCm39, using STAR. Reads quality was viewed by RseQC. Htseq-count command was used to count reads, with settings -- stranded=reverse --order=name --idattr=gene_id -- type=exon --mode=union. Package DESeq2 was used for count processing and normalization, package clusterProfiler was employed for enrichment analysis, and heatmap was plotted through heatplot.

### RNA bisulfite-seq

MYC^high^ and MYC^low^ EC4 cells (5×10^10^) were treated with either 10µM or 200µM methionine for 48h, after that total RNA was isolated with TRIzol regent. RNA quality was analyzed with Agilent RNA 6000 nano kit. 10μg total RNA was fragmented with NEBNext Magnesium RNA Fragmentation Module for 90 seconds, and samples were recovered by ethanol precipitation. DNA was removed by TURBO DNase kit. PCR template for spike-in RNA control was made from E. coli 16S RNA with NEBNext master mix, forward 5’-GAAATTAATACGACTCACTATAGGGGTGGCGGACGGGTGAGTAAT-3’, reverse 5’-TCGCACCTGAGCGTCAGTCT-3’, cycling conditions: 98°C denaturation for 10s, 62°C annealing for 30s and 72°C extension for 40s. Spike-in RNA was transcribed *in vitro* from gel purified DNA template, using MAXscrip T3/T7 Transcription Kit. Control RNA was precipitated with ammonium acetate and ethanol. RNA folding was performed on Thermo heat block, and the final spike-in control was added to samples at 1:10,000 quantity ratio. EZ RNA Methylation Kit was used to convert cytosines in RNA samples. Briefly, 1.5μg RNA samples were heated to 70°C for 10 min, followed by incubation at 64°C for 45 min, 3 circles. Samples were recovered with Zymo-Spin IC columns, diluted in 15μl nuclease free water. 3’ terminal repair was performed with NEB Quick CIP, and 5’ end phosphorylation was catalyzed with T4 polynucleotide kinase. Enzymes were removed by RNA purification kit (NEB T2030).

Seq libraries were constructed with NEBNext Multiplex Small RNA Library Prep Set for Illumina, according to the manual. Library size (~350bp) was selected using 6% polyacrylamide gel, and libraries were recovered with ethanol precipitation. After air dry for 10 min, DNA pellet was resuspended in 10μl nuclease free water. Agilent DNA 1000 kit was used to analyze library quality. Libraries were sequenced with Illumina NovaSeq 6000 system in Azenta.

Reads quality was viewed by fastqc. Cutadapt was used to filter and cut seq primers, with the parameters: -j 12 -a AGATCGGAAGAGC -A GATCGTCGGACT -e 0.1 -q 20 –M 120 --discard-untrimmed. Reference RNA was constructed with 28S rRNA (NR_003279.1), 18S rRNA (NR_003278.3), 5S RNA (NR_030686.1), 5.8S rRNA (NR_003280.2), mitochondrion (NC_005089.1) and spike_in control RNA. Reads were aligned with meRanGh, methylation rate was called with meRanCall, with settings -gref -ei 0.1 -sc 10 -fdr 0.05 -np -rl 150 -mr 0.1 -md 5 -cr 0.99 -mBQ 30 -bed63. Reads were counted with htseq-count, and size factor was estimated using estimateSizeFactors. Methylation levels between groups were compared and adjusted with the estimated size factors.

### Microarray

Microarray study (GSE73295) was performed as previously described.^29^ Briefly, LT2-MYC mice were Dox off and tumors were allowed to grow for 3 months. After that, mice were sacrificed and tumors were collected and flash frozen in liquid nitrogen. RNA was isolated, amplified and labeled with Cy3-CTP (experimental samples) or Cy5-CTP (Stratagene universal reference pool) using the Agilent low RNA input fluorescent linear amplification kits. Arrays were scanned using the Agilent microarray scanner (Agilent) and raw signal intensities were extracted with Feature Extraction v9.5 software (Agilent). Non-tumor tissues from LAP-tTTA (LT2) mice were the control.

### *In vivo* Methionine restriction study

Transgenic mice (LT2-MYC, LT2-RAS) were randomized by body weight into 4 groups (10 mice per group): MYC tumor methionine high, MYC tumor methionine low, RAS tumor methionine high, and RAS tumor methionine low. The methionine high diet (Envigo, TD180927) was a modification of diet TD.99366 to increase methionine to 8.6g/kg and remove cystine. The methionine low diet (Envigo, TD180928) was also a modification of diet TD.99366 to decrease methionine to 1.72 g/kg and remove cystine. L-alanine, L-glutamic acid and glycine was increased in methionine low diet to make the two isocaloric and isonitrogenous. Mice were maintained for 3 months. After that, livers and tumors were either fixed in 4% paraformaldehyde for histology examination or flash-frozen in liquid nitrogen.

### AZA treatment animal study

For AZA treatment, LT2-MYC transgenic mice were used. Doxycycline was removed from animal diet to initiate MYC tumor growth. After 6 weeks, mice were treated with 5 mg/kg AZA (ip, twice a week) or saline (0.9% NaCl) for another 6 weeks. 3 months later, mice were sacrificed and livers were collected for histology examination. Immunohistochemical studies were performed in a standard protocol as described in our previous work.^33^

### Nude mice xenograft study

HCC cell lines PLC/PRC/5 and HepG2 were first transfected with pLX-TRE-dCas9, selected with blasticidin for 3-5 days; secondly transfected with pDECKO containing two sgRNAs, selected for another 3-5 days. NOP2 CRISPRi sgRNA1, 5’-GCGGCCCTCCACGTGCAATC-3’; NOP2 CRISPRi sgRNA2, 5’-GCGAGTGCCGGCCGAAAGCT-3’. Non-targeting CRISPRi sgRNA1, 5’-GCTGCATGGGGCGCGAATCA-3’; Non-targeting CRISPRi sgRNA2, 5’-GTGCACCCGGCTAGGACCGG-3’.

After verification of CRISPRi knockdown, 10^10^ cells were subcutaneous inoculated in nude mice (6 weeks old). When tumor size reached 50-100 mm^3^, mice were fed with doxycycline food to induce the knockdown of NOP2. Tumor length, width and body weight were monitored every 2-3 days, until volume of most tumors in control group passed 1000mm^3^. Mice were sacrificed and tumors were removed for IHC study.

### *In vivo* Nop2 gene editing

The cloning of pDECKO-Nop2 sgRNAs were mentioned above. px330 (Addgene #327820) plasmid was double digested with XbaI and PciI, and the 8000 bp band was gel recovered. The U6-sgRNA1-H1-sgRNA2 cassette was cloned from pDECKO-Nop2 sgRNAs by PCR, and the final plasmid px330-Nop2 sgRNAs was assembled by in-fusion reactions. Sanger seq was performed to validate the final plasmid. *Nop2* sgRNAs used for *in vivo* studies: 5’-TGTCGAGTCGTGCCCGAAAG-3’ and 5’-GGAAGACGATGTGGTGACCC-3’. Non-target sgRNAs: 5’-GCGAGGTATTCGGCTCCGCG-3’ and 5’-AAATGTGAGATCAGAGTAAT-3’. Sleeping beauty plasmids pT3-EF1A-MYC-IRES-luc (Addgene #129775) and pCMV(CAT)T7-SB100 (Addgene #34879) was used to deliver MYC oncogene and luciferase into mouse liver. Hydrodynamic transfection to generate liver cancer mouse model was reported before.^34,35^ 6-week-old FVB mice were hydrodynamically injected with Sleeping beauty plasmids and px330-Nop2 sgRNAs. The ratio of pCMV(CAT)T7-SB100, pT3-EF1A-MYC-IRES-luc and px330-Nop2 sgRNAs was 0.05µg:1µg:2µg, and the plasmids were delivered in the body at 1µg/g body weight. After tail vein injection, tumor growth in the body was closely monitored by luciferase imaging. 6 weeks after hydrodynamic injection, mice were sacrificed and livers were inspected and analyzed by IHC staining.

For *Nop2* shRNA *in vivo* study, *Nop2* siRNA sequences were firstly synthesized (IDT), and cloned into pLKO.1-blast (Addgene #26655) following standard protocols. The U6-shNop2 cassette was amplified by PCR, purified with gel recovery; plasmid pT3-EF1A-MYC was double digested with NheI and PacI, then the 9000bp band was purified with gel recovery; the final pT3-U6-shNop2-EF1A was constructed by in-fusion reactions. Sleeping beauty plasmids pCMV(CAT)T7-SB100 and pT3-U6-shNop2-EF1A-MYC were injected at a ratio of 0.05µg:1µg, 1µg/g body weight into mouse liver. *Nop2* siRNA (SI01328992, Qiagen) was: 5’-AAGACGAACAAGGATGAGAAA-3’, scramble (1027280, Qiagen): 5’-CAGGGTATCGACGATTACAAA-3’.

## QUANTIFICATION AND STATISTICAL ANALYSIS

Results were mean ± SEM of three independent experiments unless otherwise noted. Unpaired Student’s t-test and Spearman correlation coefficient analysis were used. P values were two-sided: 0.05 was considered statistically significant.

## Results

### Methionine metabolism in MYC-driven liver cancer

We explored metabolic vulnerabilities in MYC-driven liver cancer using LAP-tTA×TetO-MYC (LT2-MYC) transgenic mice, a conditional model that drives MYC-dependent tumorigenesis in the liver and has features of human HCC and hepatoblastoma^29,36^ (Figure 1A, Figure S1A). We performed mass spectrometry-based steady state metabolite analysis of LT2-MYC tumors compared to normal livers.^15^ We found that methionine as well as multiple metabolites of the methionine metabolism pathway were significantly altered in MYC-driven liver cancer (Figure 1B, E, Figure S1B-S1F), with most methionine-related metabolites higher in MYC tumors (Figure 1B, E and Figure S1C) compared to control livers. To determine whether alterations in the methionine pathway is MYC oncogene specific, we compared metabolomic data from liver tumors driven by an activated RAS oncogene (LT2-HRAS^G12V^).^29^ We found that abundance of methionine associated metabolites was enriched in MYC tumors compared to RAS tumors (Figure S1D-S1F), suggesting MYC may differentially regulate methionine metabolism.

**Figure 1.**
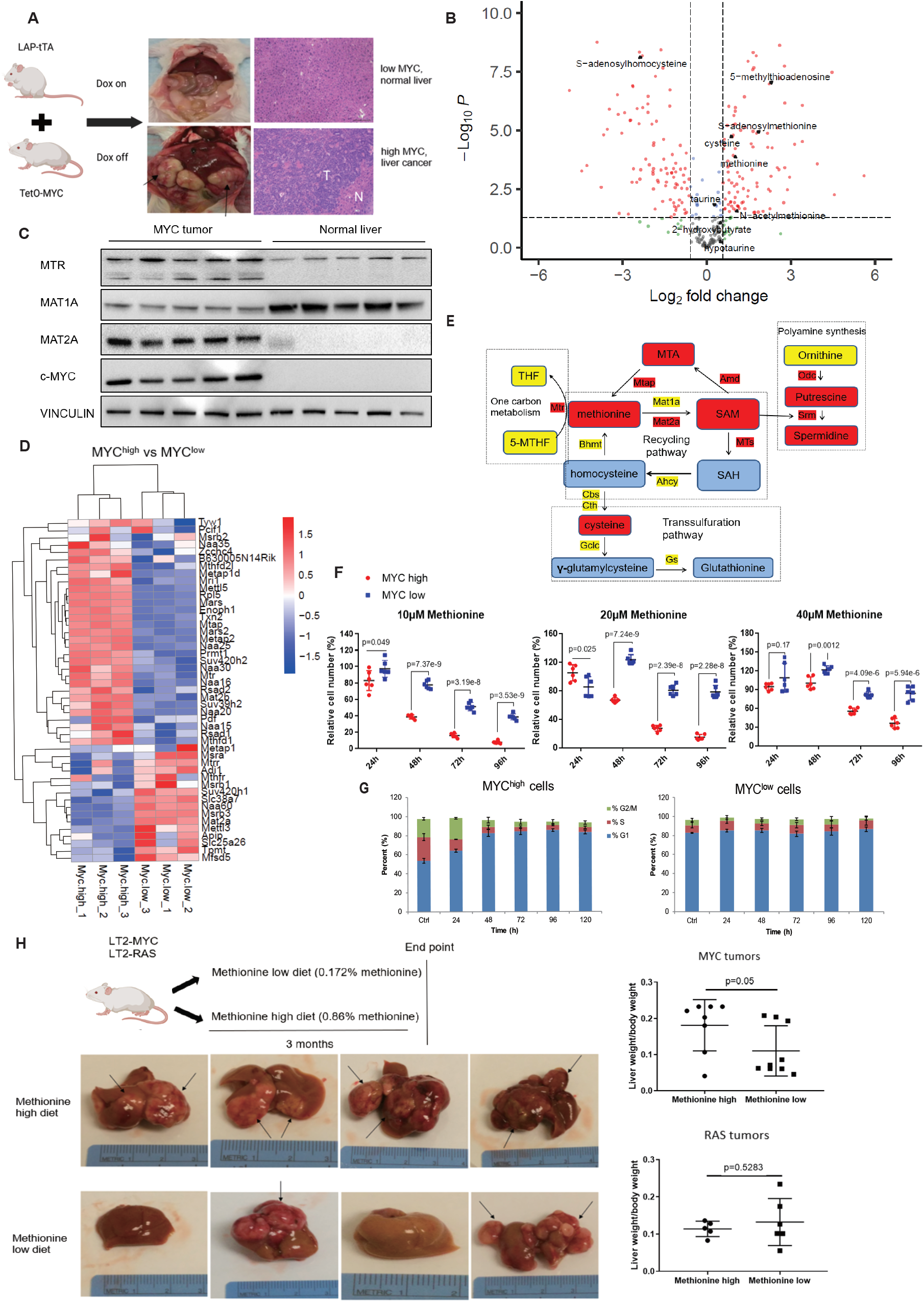
Altered methionine metabolism in MYC-driven liver cancer. (A) LT2-MYC liver cancer transgenic model in which doxycycline is removed from the diet to induce liver tumor formation. Tumor is labeled as T and adjacent tissue is N. (B) Volcano plot for metabolites in MYC tumors versus normal livers. Methionine-pathway related metabolites were shown. (C) Protein levels of methionine metabolism enzymes Mtr, Mat1a, Mat2a as well as Myc in normal livers and MYC-driven tumors. (D) Heatmap of methionine related genes in EC4 MYC^high^ vs MYC^low^ cells (RNA seq). (E) Schema depicting methionine metabolism pathways. Metabolites and enzymes are compared between MYC-driven liver cancer and normal livers. Data were sourced from metabolites (rectangles) and microarray (without box). Red, significantly increased at p≤0.1; blue, significantly decreased at p≤0.1; yellow, no significant change detected. (F) Low methionine medium significantly inhibits MYC^high^ cell proliferation. MYC^high^ and MYC^low^ EC4 cells were cultured with 10-40 µM methionine or control full media (200 µM) for 24-96 h; relative cell number was calculated by cell counting viable cells. (G) Methionine starvation induced increased G1 cell cycle predominantly in MYC^high^ cells. FACS DNA content analysis showing proportion of cells in each phase over time for MYC^high^ (left) and MYC^low^ cells (right). (H) Low methionine diet inhibited MYC tumor growth *in vivo*. Transgenic mice bearing MYC or RAS tumors were fed with either low (0.172%) or high methionine (0.86%) diet for 3 months, after that mice were sacrificed. Left: photos of MYC tumors were shown (methionine high vs low). Arrows indicate tumors. Right: ratio of liver+tumor weight versus total body weight. Only in MYC tumors, the ratio was diminished in methionine low diet.

Methionine is regenerated from homocysteine; the reaction is catalyzed by methionine synthase (MTR). Methionine is catalyzed by methionine adenosyltransferases (MATs) to produce S-adenosyl-methionine (SAM, or AdoMet), the active form of methionine used by methyltransferases. Downregulation of MAT1A and upregulation of MAT2A, known as MAT1A:MAT2A switch, is associated with hepatocellular carcinoma (HCC).^37^ We saw altered expression of methionine metabolism enzymes proteins in MYC-driven liver cancer compared to normal liver, including increased levels of MAT2A and MTR, and decreased levels of MAT1A (Figure 1C), which was in accordance with the MAT1A:MAT2A switch.

To determine if changes in methionine-pathway genes are temporally regulated by MYC expression, we used a conditional murine liver cancer cell line, EC4, which was derived from the LT2-MYC tumors.^29^ MYC expression can be inhibited in the cells with doxycycline treatment allowing us to determine acute changes in gene expression upon MYC inhibition.^15^ Cysteine and methionine KEGG pathway genes were significantly enriched in MYC^high^ cells (Figure S1G and S1H). We generated a list of methionine-related genes from gene ontology (www.geneontology.org). Compared with MYC^low^ cells, gene expression of most methionine-related genes was increased in MYC^high^ cells (Figure 1D). We next examined methionine related genes in MYC-driven tumors versus normal livers. Similarly, many methionine related genes were also upregulated in MYC tumors (Figure S1I). We next integrated RNA-seq data (MYC tumors), steady-state metabolite data, and enzyme protein expression (western blot) in MYC-driven liver cancer (Figure 1E). We observed a high SAM:SAH ratio in MYC-driven liver tumors (2.70±0.72) compared to control livers (0.15±0.04). Prior studies have suggested that a high SAM:SAH ratio correlates with hypermethylation,^37^ suggesting that methyltransferase activity may be increased in MYC tumors.

We next sought to determine whether MYC-dependent tumor growth requires abundant methionine. Methionine concentration varies from 5 to 40 µM in the plasma of healthy individuals;^38^ however, typical cell culture medium (i.e. DMEM) contains much more methionine (200µM). When EC4 cells were cultured in physiological methionine conditions (10-40 µM) MYC^high^ cell growth was more potently inhibited than MYC^low^ cells at lower methionine concentrations compared to full media (methionine 200µM) (Figure 1F). To determine whether the same methionine dependence is observed in human liver cancer cells, we examined a panel of five human HCC cell lines with varying MYC protein expression levels. Cell growth in 5 µM methionine was dramatically inhibited in Hep40 and SNU398 cells that expressed high levels of MYC. In contrast, SNU475 cells that express low MYC were much less inhibited (Figure S1J). We next asked if partial depletion of MYC with siRNA, could reverse dependence on methionine. Knockdown of MYC with a mixture of 4 siRNAs greatly alleviated liver tumor cell growth dependence on methionine (Figure S1K) suggesting that methionine is especially important for the growth of MYC high HCC cells. Growth under low methionine conditions induced G1 cell cycle arrest only in EC4 MYC^high^ cells (Figure 1G and Figure S1L), with G1 phase increased from 53.53±2.55% (Ctrl) to 83.47±2.06% (at 120 h, p=0.000125). While MYC low cells have a higher proportion of cells in G1 at baseline, this was not significantly increased in low methionine conditions (Figure 1G).

The most stringent test of methionine dependence is to determine if depleting methionine *in vivo* alters *de novo* tumor formation. We evaluated the effect of dietary methionine on tumor growth in the transgenic MYC-driven (LT2-MYC) or RAS-driven (LT2-RAS)^29^ models of liver cancer by using diets containing different concentrations of methionine. Diets were isocaloric and isonitrogenous, except for the methionine (0.172% in low methionine diet versus 0.86% in high methionine group), were used for these studies. Mice were removed from a doxycycline diet to induce oncogene expression and were fed either a methionine high or low diet for 3 months, then mice were sacrificed, and tumors were analyzed. In the MYC group, mice fed with low methionine diet developed smaller tumors, compared with those given a methionine high diet (Figure 1H). In contrast, a low methionine diet did not affect the growth of RAS tumors, indicating only MYC tumors were dependent on methionine.

### MYC tumors are depended on methylation

We next determined if limiting methionine would alter MYC-driven transcription. MYC^high^ and MYC^low^ cells were cultured with either 10 µM or 200 µM methionine medium for 24 h, and RNA was isolated for sequencing. In the presence of standard methionine (200 µM), toggling MYC ON/OFF induced significant transcriptional changes, including MYC increased gene expression related to ribosome and ncRNA processing (Figure 2A). Interestingly, ribosome and ncRNA gene sets were among the top enriched programs in methionine high vs low conditions (Figure 2B). Since both MYC overexpression and high methionine conditions regulated similar gene sets, we asked if methionine abundance could change MYC expression. Both MYC protein and MYC S62 phosphorylation, which increased MYC transcriptional activity,^39^ were increased in a methionine dependent manner in both Hep40 and SNU398 cells (Figure 2C). Even in EC4 cells where MYC protein expression was driven by an exogenous promoter we also observed methionine regulated MYC expression (Figure S2A), suggesting effects may be post-transcriptional or related to the proliferative state of the cells. These results suggest that by depleting methionine we might be able to attenuate MYC protein abundance and hence function.

**Figure 2.**
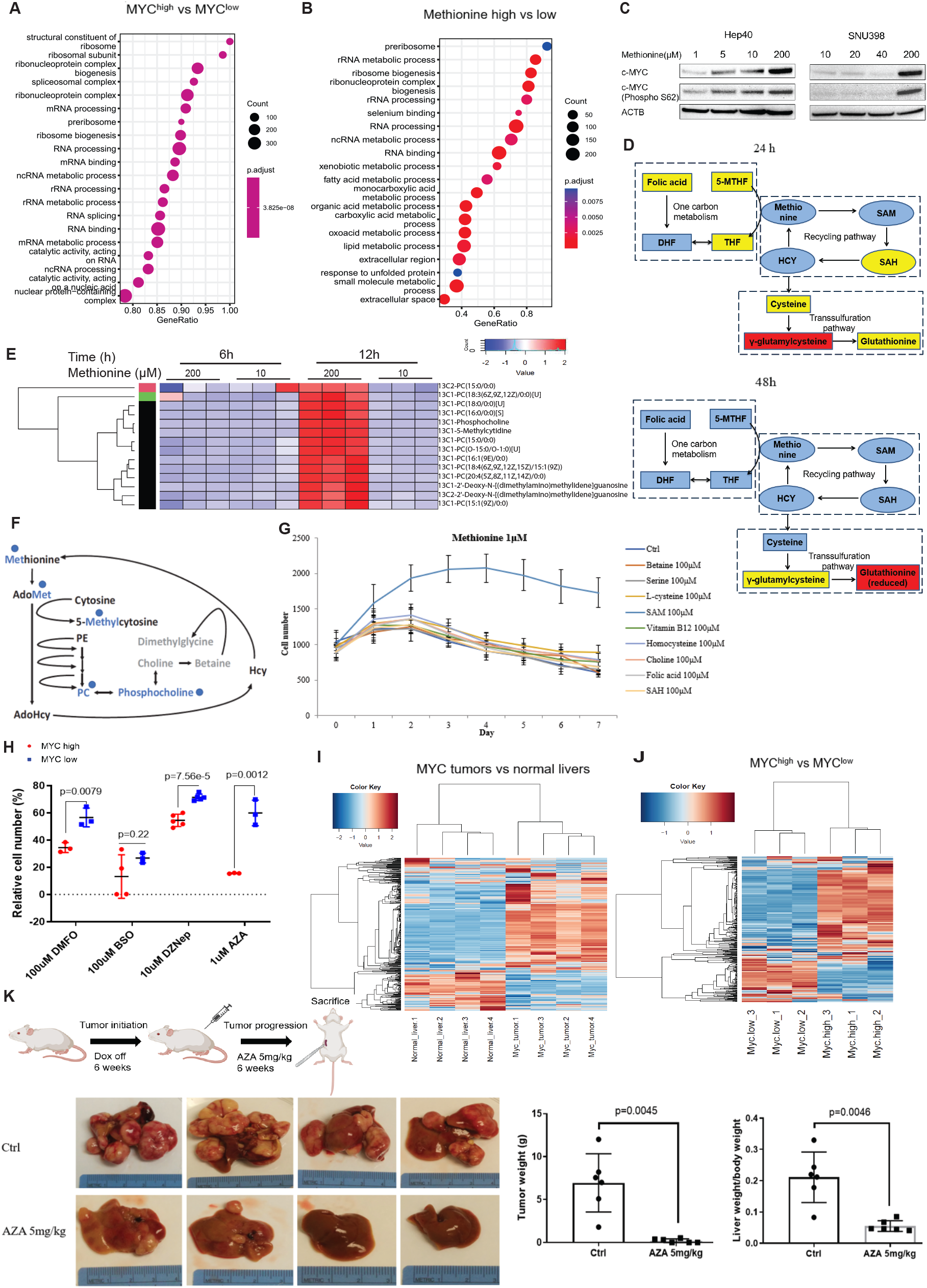
MYC-driven liver tumorigenesis is depended on methylation. (A) GO enrichment of all the differentially expressed genes (DEGs) in conditional EC4 cells, MYC^high^ vs MYC^low^ (RNA seq). Cells were grown in 200 µM methionine. (B) GO enrichment of all the DEGs in conditional EC4 cells, methionine high (200µM) vs low (10µM) when MYC expression is induced (MYC^high^) (RNA seq). (C) Western blot showing MYC expression and MYC S62 phosphorylation. Actin (ACTB) as loading control. (D) Relative abundance of significantly altered methionine pathway metabolites, MYC^high^ vs MYC^low^. Cells cultured with 20 µM methionine for 24 and 48 h, after that cell number was counted, and cell samples were collected for LC-MS detection. The final results were adjusted with cell number. Red, significantly increased in MYC^high^ cells at p ≤ 0.05; blue, significantly decreased in MYC^high^ cells at p ≤ 0.05; yellow, no significant change. (E) 13C-methionine tracing in EC4 MYC^high^ cells. Cells were washed with PBS to remove media, then cultured either with 200µM or 10µM methyl-13C methionine for 6h or 12h, and finally subjected for mass spectrometry. Alterations of 13C labeled metabolites were shown in heatmap. (F) Schematic diagram of 13C-methionine tracing. 13C labeled methyl group was depicted in blue and shown as a dot. AdoMet is S-adenosyl-L-methionine, or SAM. (G) SAM partially rescued MYC^high^ cells in low methionine medium. Cells were cultured with DMEM containing 1µM methionine for 7 days, with or without 100µM betaine, serine, L-cysteine, SAM, vitamin B12, homocysteine, choline, folic acid or SAH, respectively. (H) MYC^high^ cells were most sensitive to the inhibition of DNA/RNA methylation. MYC^high^ and MYC^low^ EC4 cells were treated with inhibitors targeting DNA, RNA, protein methylation and SAM downstream pathways for 72 h, and cell proliferation was analyzed. DMFO, inhibitor of polyamine synthesis pathway. BSO, inhibitor of transsulfuration pathway. DZNep, histone methylation inhibitor. AZA, nucleotide methylation inhibitor. (I) Clustering for methylation genes in MYC tumors vs normal livers. Methylation genes were sourced from Gene Ontology Resource (https://geneontology.org/). (J) Clustering for methylation genes in EC4 cells, MYC^high^ vs MYC^low^ (K) AZA treatment inhibited MYC tumor growth. MYC expression was induced to allow for the initiation of tumor formation for 6 weeks, after that mice were administrated 5 mg/kg AZA (ip, twice per week) for another 6 weeks. After 3 months mice were sacrificed, and macroscopic tumors were dissected and weighed. Saline served as the control (n=6). Left: Representative images of livers at treatment endpoint. Middle: Quantification of liver tumor weight middle panel for control and AZA treated groups. Right: ratio liver+tumor weight compared to total body weight.

To understand the mechanism of methionine dependence in MYC-driven liver cancer, we compared metabolites in switchable MYC^high^ and MYC^low^ EC4 cells (Figure 2D). Cells were cultured in 20 µM methionine (mild restriction) for 24 h or 48 h, and metabolomics analysis was performed by LC-MS. At 24 h, changes were mostly restricted to the methionine recycling pathway (Figure S2B); by 48 h, however, nearly all the methionine pathway metabolites were markedly decreased in MYC^high^ cells, except for glutathione (reduced) (Figure S2C).

The methionine recycling pathway generates SAM, the dominant methyl donor for methylation reactions. We used methyl-13C methionine to trace the methyl group distribution in the EC4 cells that were cultured either with 200µM (high) or 10µM (low) methyl-13C methionine for 6h or 12h followed by mass spectrometry. At 6h, there was no difference on SAM levels between methionine high or low; however, at 12 h SAM levels significantly increased in methionine high but depleted in methionine low medium (Figure S2D). We found 13C labeled 5-methylcytidine, phospholipids and 2’-Deoxy-N-[(dimethylamino)methylidene]guanosine were much more abundant in methionine high medium than in methionine low conditions (Figure 2E). 2’-Deoxy-N-[(dimethylamino)methylidene]guanosine is a precursor for nucleotide synthesis. Phospholipids contribute to cell membrane biosynthesis and can be used for signal transduction. 5-methylcytidine can be derived from 5C-methylated-RNA or 5C-methylated DNA modified. Thus, the tracing studies suggest methionine-derived methylation contributes to RNA and/or DNA m5C methylation, cell membrane synthesis and DNA synthesis (Figure 2F).

To further define which methionine-derived metabolites are essential for the proliferation of MYC^high^ cells, we conducted metabolite rescue assays. Methionine-related metabolites such as SAM, SAH, folic acid, choline, homocysteine, cysteine, serine, betaine and vitamin B12 were added back to media which contained only 1 µM methionine and the proliferation of MYC^high^ cells was examined (Figure 2G). While none of the metabolites could fully rescue methionine depletion, SAM was the only metabolite able to partially rescue the proliferation of MYC^high^ cells over the course of one week (Figure 2G). We next examined how protein expression of key methionine metabolism mediating enzymes was altered in the context of methionine depletion by western blot (Figure S2E). All the enzymes had higher levels in MYC^high^ cells when cells were grown in full methionine medium. However, under methionine depleted conditions, only enzymes that produce SAM, specifically MTR, MAT1A, and MAT2A, still showed higher levels in MYC^high^ cells (Figure S2E). Following methionine depletion enzymes that consume SAM, including MTAP, CBS and SHMT2, a key enzyme of one carbon metabolism, all showed lower expression. These data suggest that under restricted methionine conditions, MYC^high^ cells downregulate SAM utilization, indicating an essential role for SAM in MYC^high^ cells.

SAM is essential for both DNA, RNA, protein methylation and downstream pathways like polyamine synthesis and transsulfuration pathway. To further investigate to which MYC high cells are most sensitive, EC4 cells were treated with inhibitors that specifically target methylation reactions and downstream pathways, and cell viability was measured. Amongst the various methionine pathway perturbations, we observed that MYC^high^ cells were most sensitive to AZA, an inhibitor of DNA and RNA methylation (Figure 2H).^40^ AZA dramatically inhibited MYC^high^ cell proliferation in a dose dependent manner, whereas its effect on MYC^low^ cells was much less (Figure S2F). These findings are consistent with the results of the methionine tracing study, which showed the methyl group was incorporated into ribonucleotides and deoxynucleotides.

Given the ability of the methylation inhibitor AZA to attenuate proliferation of MYC^high^ EC4 cells, we examined which methylation-associated genes are altered in a MYC-dependent manner in EC4 cells. Many methylation related genes were highly transcriptionally expressed in MYC tumors and Myc^high^ cells (Figure 2I and 2J). These included methylation related gene sets, like methyltransferase complex, DNA methylation, macromolecular methylation and RNA methyltransferase complex, which were significantly enriched (Figure S2G). We also observed that genes associated with nucleotide modifications including methylation were enriched in MYC tumors and MYC^high^ cells (Figure S3A-S3C).

Given the dependence on SAM and alteration in methyltransferase pathway genes in MYC tumors, we asked if inhibition of DNA/RNA methylation could affect tumor formation. Azacitidine (AZA) is an approved drug for myelodysplastic syndrome, and is also used for myeloid leukemia, and juvenile myelomonocytic leukemia.^41^ AZA was reported to inhibit both DNA and RNA methylation.^42,43^ To test its effects in MYC-driven liver cancers, LT2-MYC transgenic tumors were initiated by removal of doxycycline from mouse diet for 6 weeks. Mice were then treated with 5 mg/kg AZA (intraperitoneal injections, twice per week) or diluent for 6 weeks. We found that AZA treatment potently inhibited MYC tumor growth, with only rare tumor nodules noted at the time of tumor collection and a significant decrease in liver/total body weight (Figure 2K). Thus, MYC-driven tumorigenesis was markedly inhibited by AZA, a broad inhibitor of DNA/RNA methylation.

### NOP2 is a MYC transcriptional target

Given the importance of SAM for MYC tumor cell proliferation and the ability of AZA to markedly diminish tumor formation, we reasoned that one or more RNA or DNA methyltransferases were important for MYC-driven tumor growth. Analysis of RNA seq data identified 39 SAM-dependent methyltransferases (MTs) significantly changed in MYC tumors compared to normal liver, while the expression of 71 SAM-dependent MTs were altered in EC4 cells when MYC was toggled ON / OFF (Figure S4A). A total of 26 MTs were identified to be upregulated in both MYC tumors and MYC^high^ cells. Of these, five m5C methyltransferases were found: including rRNA methyltransferase *Nop2* and *Nsun2*, and DNA methyltransferase *Dnmt1, Dnmt3a* and *Dnmt3b* (Figure 3A).

**Figure 3.**
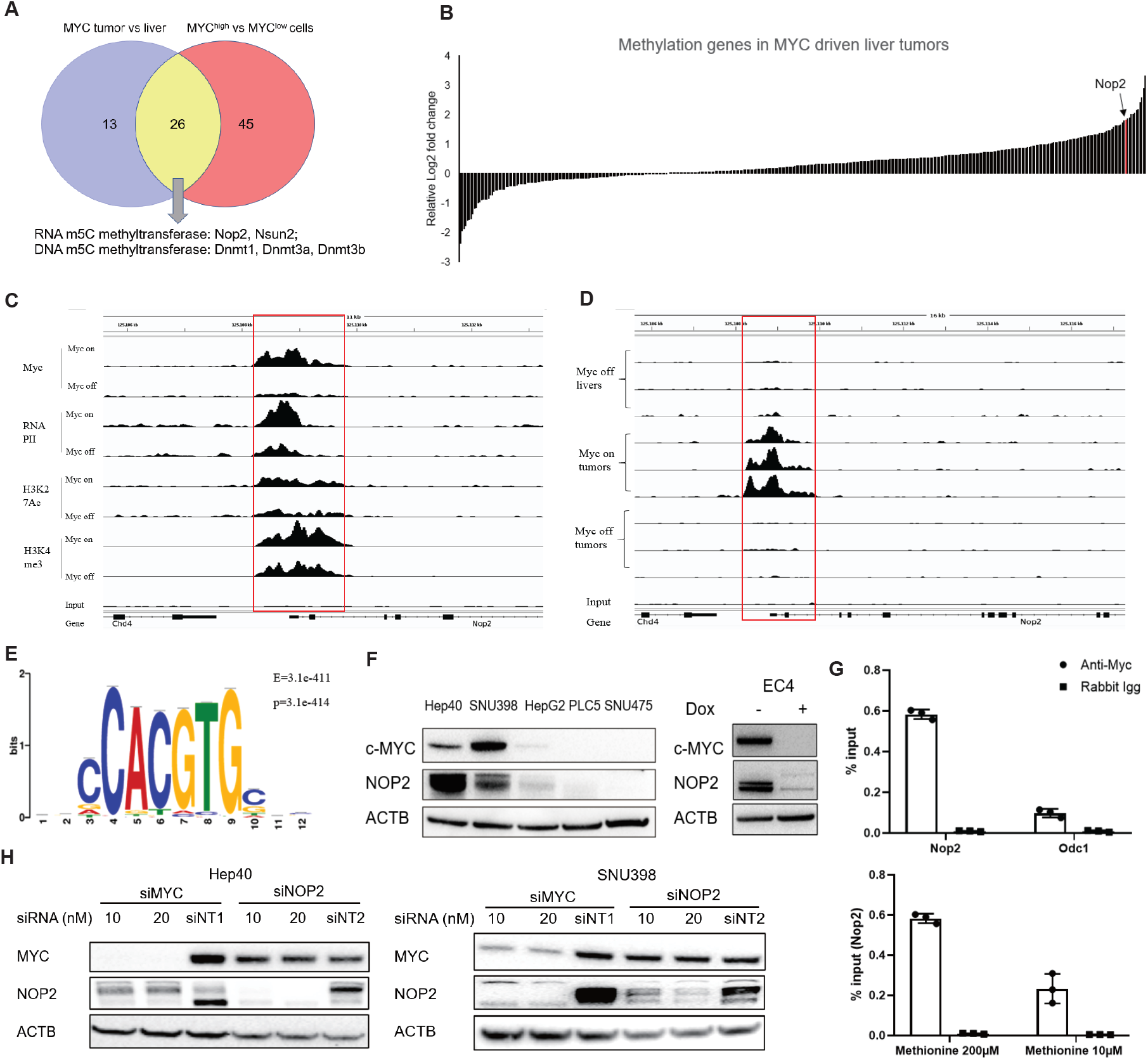
RNA methyltransferase NOP2 is a MYC transcriptional target. (A) Upregulation of SAM dependent methyltransferases in MYC tumors (tumors vs normal livers) and MYC^high^ EC4 cells (MYC^high^ vs MYC^low^). Five m5C methyltransferases were found significantly increased in both MYC tumors and MYC^high^ cells. (B) Gene expression (Log2 fold change) of methylation-associated genes in MYC tumors compared with normal livers (n=4). The fold-increase of Nop2 mRNA expression was one of the highest. (C) The binding of Myc, RNAP II (RNA polymerase II), H3K27Ac and H3K4me3 on the Nop2 promoter (GSE83869, Myc ON vs OFF; visualized with the IGV browser). (D) Status of Myc expression on the Nop2 promoter, Myc OFF liver vs Myc ON tumor vs Myc OFF tumor (GSE76078, IGV browser). (E) Canonical MYC binding site, E-box, was found in Nop2 core promoter. MEME ChIP was used for DNA motif analysis. (F) NOP2 and MYC protein levels in human HCC cell lines and MYC^high^, MYC^low^ EC4 cells. (G) ChIP assay in EC4 cells. Cells were cultured in either 200µM or 10µM methionine media for 24h. MYC target gene Odc1 served as the positive control, and purified rabbit IgG was used as the negative control. (H) Silencing of MYC in SNU398 and Hep40 cells decreased NOP2 protein expression, while knockdown of NOP2 did not affect MYC levels. Cells were treated with siRNAs (Dharmacon ON-TARGETplus SMARTPool) specifically designed against MYC or NOP2 for 72h, and western blot was performed.

Changes of MTs *in vivo* could be a direct or indirect effect of MYC-driven tumorigenesis. A recent study examined the LT2-MYC mice in which MYC expression *in vivo* is induced for only 48 hours then mice were sacrificed and livers were subjected to RNA seq and ChIP seq analysis (GSE83869).^44^ Of these 26 MTs, expression of *Trmt11, Nop2* and *Prmt3* was significantly increased after MYC was acutely induced *in vivo* (Figure S4B), suggesting they are direct transcriptional targets of MYC *in vivo*. Only *Nop2* has m5C methyltransferase activity. NOP2 has been found to methylate human 28S rRNA at C4447,^45^ and this modification is crucial for ribosome biogenesis^46^ and efficient function. NOP2 also promotes cell proliferation and tumor progression.^47,48^ Ribosome related genes were enriched in MYC tumors and MYC^high^ cells (Figure 2A and 2B, Figure S4C and S4D), and among these a MYC-dependent increase in *Nop2* expression was one of the highest (Figure 3B).

To explore whether *Nop2* transcription is regulated by MYC, we analyzed MYC ChIP seq data (GSE83869). MYC induction in livers for 48 h greatly increased MYC binding to the transcriptional start sites (TSS) of many genes, however the binding of RNA polymerase II (RNAP II), H3K27Ac and H3K4me3, changes associated with active transcription, did not significantly change for most MYC-bound genes after only 48 h of MYC induction (Figure S5A). However, there were several exceptions, indication early transcriptional effects of MYC binding. For example, MYC binding was enriched at ribosome and ncRNA gene promoters (Figure S5B). MYC specifically bound to *Nop2* promoter; together with RNAP II, suggesting direct transcriptional activation of Nop2 (Figure 3C). *Odc1* is a well-established MYC transcriptional target gene and we saw both MYC and RNAP II binding increased on *Odc1* promoter (Figure S5C). The increase of MYC or RNAP II binding to the TSS was not observed in the other methyltransferases (Figure S5C). Finally, MYC binding to the *Nop2* promoter was confirmed with another CHIP seq dataset GSE76078,^49^ which also used conditional LT2-MYC mice. MYC binding was significantly increased in MYC tumors, and this binding reversed after MYC expression was turned off (dox on for 16 h) (Figure 3D). DNA motif analysis identified the canonical E-box, CACGTG, in *Nop2* core promoter (chr6:125108894-125108905(-); Figure 3E). We examine how NOP2 protein expression corelated with MYC expression and found that NOP2 protein was abundant in MYC high cell lines Hep40 and SNU398 and EC4 MYC^high^ cells but expressed at lower levels in cells with diminished MYC expression (Figure 3F). MYC binding was validated with ChIP assay in EC4 cells, demonstrating the enrichment of MYC on the *Nop2* promoter, but diminished binding under low methionine (10 µM) conditions (Figure 3G). Depletion of MYC with a pool of siRNAs in Hep40 and SNU398 cells also decreased NOP2 protein, but knockdown of NOP2 did not affect MYC levels (Figure 3H). *NOP2* mRNA levels (RNA seq data) correlated with *MYC* mRNA expression (Figure S5D), MYC target gene sets (including hallmark MYC targets) (Figure S5E) and MYC oncogenic signature or active pathway (Figure S5F) in dozens of liver cancer cell lines from the Board Institute’s Cancer Dependency Map (DepMAP) project. Together, these data demonstrate that *NOP2* is a direct MYC transcriptional target gene and *NOP2* expression can be diminished with MYC depletion.

### NOP2 associates with MYC expression and worse patient outcome

We explored NOP2 and MYC expression association in liver cancer clinical datasets. Among a cohort of 360 liver cancer samples from TCGA, we divided the samples into 2 groups, MYC mRNA high (MYC mRNA >83.06%, n=61) and low (MYC mRNA≤83.06%, n=299) based on an optimized cut-off. Gene set enrichment analysis (GSEA) showed that ribosome biogenesis, rRNA processing, translation etc. were top enriched in MYC high patients (Figure S6A and S6B). MYC gene expression modestly correlated with RNA methylation gene sets, but did not relate to DNA methylation pathways, histone methylation and protein methylation pathways (Figure 4A). There are multiple types of RNA methylation. Except for m5C methyltransferase *NOP2*, we also found the fold change of 2’-O-methyltransferase *FBL* was the highest in MYC tumors (Figure S4A). 2′-O-methylation is the most abundant rRNA modification in ribosomes. Recent research demonstrated that FBL was essential for tumorigenesis,^50^ so *FBL* was also included in the analysis. We analyzed the five m5C methyltransferases (*NOP2, NSUN2, DNMT1, DNMT3A, DNMT3B*) shown as a volcano plot (*MYC* high patients vs MYC low patients). *NOP2* was the most significantly up-regulated gene (Figure 4B). We analyzed the expression of *MYC* and the five m5C methyltransferases and *FBL* in TCGA samples, and found only *NOP2* mRNA modestly correlated to *MYC* gene expression, with a correlation efficiency of 0.3277 (p=1.97e-10) (Figure 4C). *NOP2* mRNA was significantly higher in *MYC* high cohort (Figure 4D). Pathway analysis demonstrated that *NOP2* expression within liver cancer TCGA dataset associated with cell cycle, DNA replication and DNA repair, chromatin remodeling, histone and DNA methylation, etc (Figure 4E and S6C). Similarly, we found *MYC* was highly associate with NOP2 (R=0.48, p=3.6e-15), while modestly correlated to *FBL* (R=0.24, p=1.8e-4) and *DNMT3A* (R=0.28, p=1.5e-5) in CLCA dataset (China Liver Cancer Atlas, n=239) (Figure S6D).^51^ Ribosome biogenesis, rRNA processing, ribonucleoprotein complex biogenesis, etc. were highly enriched in *MYC* high group (CLCA, n=34) when compared with *MYC* low cohort (CLCA, n=205) (Figure S6E). ssGSEA analysis showed NOP2 expression was also significantly associated with MYC signature and MYC targets, including hallmark MYC targets (CLCA, Figure S6F).

**Figure 4.**
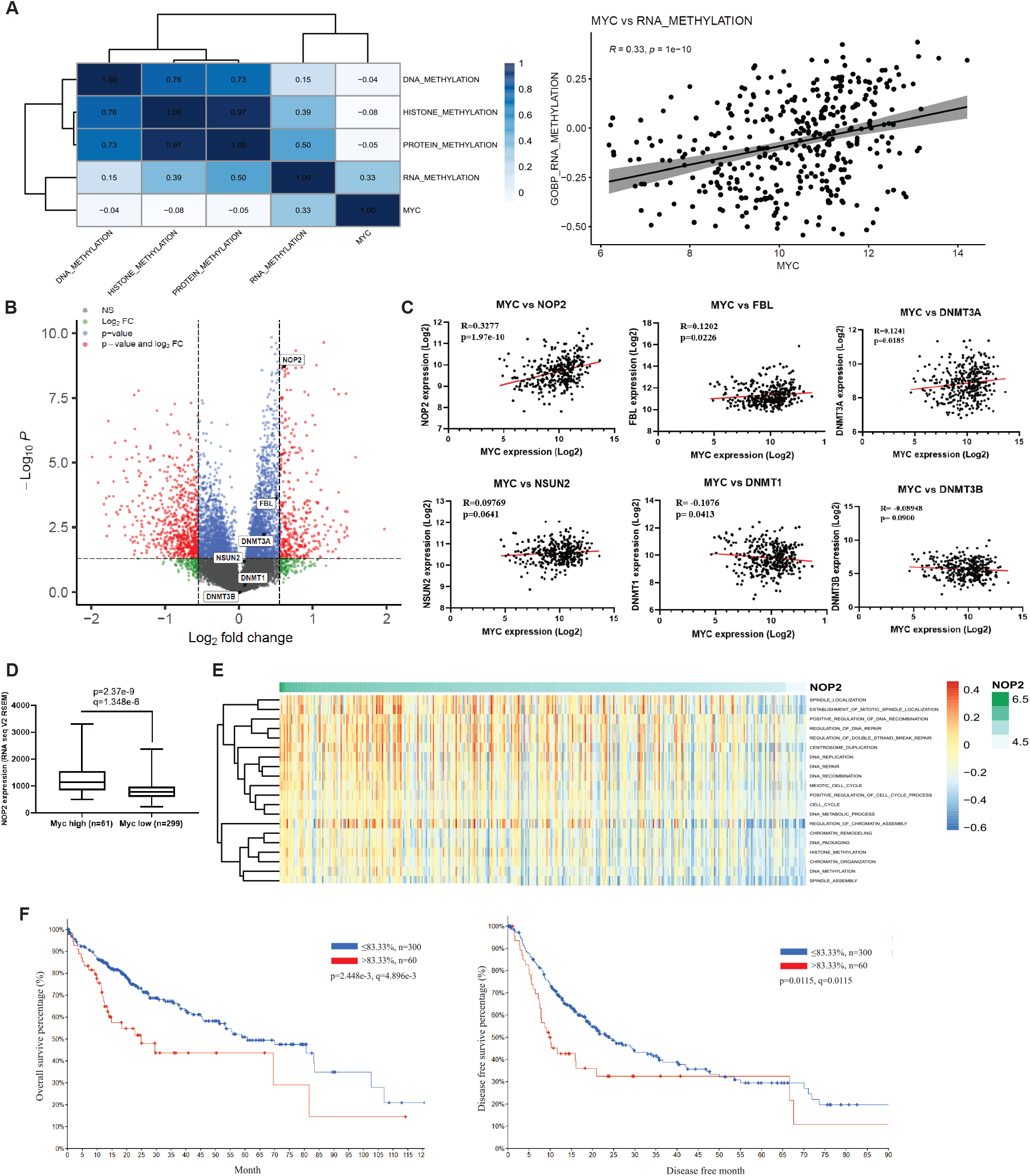
NOP2 associates with MYC expression and worse patient outcome. (A) Correlation analysis between MYC gene expression and DNA, RNA, protein, histone methylation in TCGA liver cohort. (B) Volcano plot of TCGA clinical data, MYC high versus MYC low. Five m5C methyltransferases as well as FBL was shown. (C) NOP2 expression modestly correlated with MYC mRNA levels in TCGA cohort (n=366, R=0.3277, p=1.97e-10). (D) NOP2 levels were significantly high in MYC high patients, TCGA cohort. (E) Heatmap of NOP2 associated biological pathways in TCGA cohort. NOP2 expression associated with cell cycle, DNA replication, centrosome duplication, chromatin organization, DNA and histone methylation in the clinic. (F) NOP2 expression associated with poor clinical outcomes (overall survival and disease-free survival, TCGA cohort).

*NOP2* was also associated with poor outcome features including: elevated *AFP* tumor marker levels (Figure S6G), higher liver tumor grade (Figure S6H), and more vascular invasion (Figure S6I) compared to *NOP2* low tumor samples. Elevated *NOP2* expression is associated with poor prognosis, as demonstrated by overall survival and disease-free survival (Figure 4F) in the TCGA cohort. We also explored *NOP2* expression across a variety of human cancers, and found *NOP2* was highly expressed in other tumors (Figure S6J) and associated with poor survival (Figure S6K). Collectively, these demonstrated that *NOP2* associates with MYC and poor clinical outcomes.

### NOP2 methyltransferase activity and methionine dependence in MYC-driven liver cancer cells

We next sought to determine how NOP2 m5C rRNA methyltransferase activity is altered in the context of different methionine concentrations. We performed bisulfite sequencing in conditional MYC EC4 cells to determine rRNA (28S, 18S, 5S and 5.8S) methylation. Only 28S rRNA was found to be methylated. The methylation rate at 3 cytosines were significantly lowered when we compared to methionine high (200µM) vs low (10µM); these included C3438, C3683 and C4099 of 28S (Figure 5A). The sequences of the three positions were also shown to the left. The sequence of C4099 in mouse EC4 cells corresponds to C4447 in human 28S rRNA, which is the previously established methylation site for NOP2. Compared with 200 µM, 10 µM methionine reduced C4099 methylation, which clearly demonstrated the effects of limiting cellular access to methionine. Knockdown of *Nop2* with a pool of siRNAs also reduced C4099 methylation rate similar to culturing in 10 µM methionine (Figure 5B), further confirming a role of NOP2 in regulating 28S rRNA. The *Nop2* siRNA pool also slightly reduced C3438 methylation but did not alter C3683 methylation. However, low methionine-media did not significantly decrease C4099 methylation in MYC^low^ EC4 cells, suggesting that methylation at this site was dependent on a MYC-methionine-NOP2 axis (Figure 5C).

**Figure 5.**
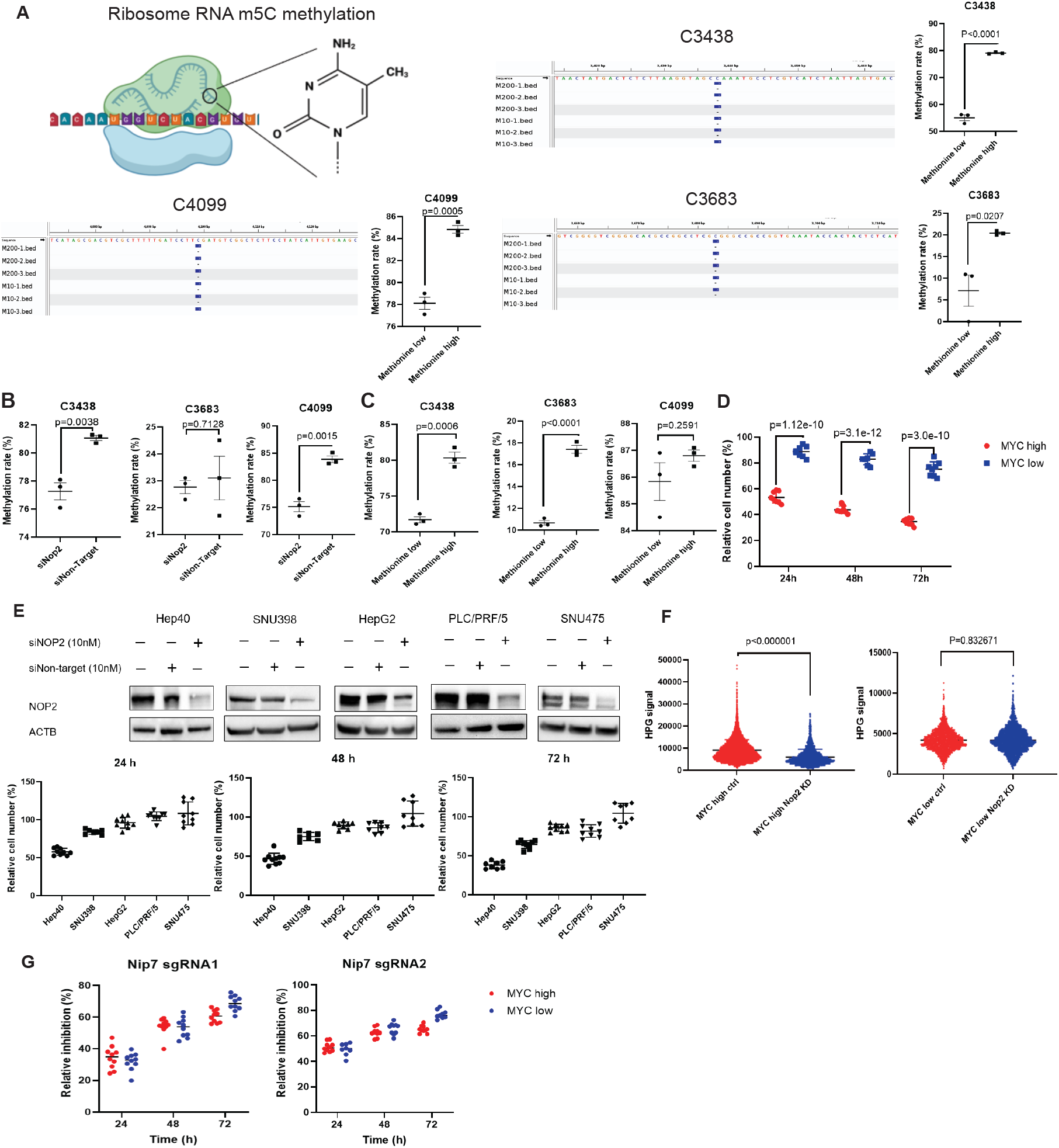
NOP2 is involved in the methionine dependence of MYC-driven liver cancer cells. (A) m5C methylation levels in 28S rRNA, methionine high (200µM) vs low (10µM) in MYC^high^ cells. Bed files were shown (IGV browser). (B) Methylation levels in 28S rRNA at sites C3438, C3683, and C4099 following treatment with siNop2 versus siNon-target control. (C) m5C methylation levels in 28S rRNA, methionine high (200µM) vs low (10µM) in MYC^low^ cells. (D) MYC^high^ cells were more potently inhibited by Nop2 CRISPR knockdown (Nop2 sgRNA6). Cells were infected with Nop2 sgRNA6 virus for 48h, after that cells were selected with puromycin for up to one week. Infected cells were used for cell growth assay, and cell proliferation was determined by cell number counting. Non-target sgRNA was used as control. (E) Depletion of NOP2 in HCC cell lines. MYC high cell lines Hep40 and SNU398 were much more sensitive to NOP2 inhibition. (F) Fluorescence quantification between MYC^high^ Nop2 KD cells and normal MYC^high^ cells, and between MYC^low^ Nop2 KD cells and Ctrl MYC^low^ cells. Nascent protein synthesis was measured by Click-iT HPG kit. (G) Knockdown of Nip7 significantly reduced both MYC^high^ and MYC^low^ EC4 cell proliferation. Cells were transfected with Nip7 sgRNAs for 48h, followed by a puromycin selection for 3 days. Cell proliferation was assayed by counting cell numbers. Non-target sgRNA was used as control.

We next asked whether NOP2 expression is required for liver cancer cell proliferation. We generated multiple *Nop2* sgRNAs and found that sgRNA6 worked best (Figure S7A) in a transient assay to deplete NOP2. EC4 cells expressing Cas9 were infected with *Nop2* sgRNA6 virus, selected with puromycin for one week and cell proliferation was measured. Compared with MYC^low^ cells, MYC^high^ cells were much more potently inhibited after infection with sgRNA virus (Figure 5D). We also measured the effect of *NOP2* silencing on HCC cell proliferation. Similarly, siRNA knockdown of *NOP2* greatly diminished the growth of MYC-high human liver cancer cell lines, Hep40 and SNU398, more than that of MYC-low cells, such as SNU475 (Figure 5E). We also tried to generate a *Nop2* knockout EC4 cell lines with CRISPR technique; however, EC4 cells couldn’t survive after *Nop2* loss (Figure S7B and S7C), indicating *Nop2* is an essential gene for cancer cell growth. There might be counterselection for complete loss of Nop2 but we finally got a knockdown (KD) clone, by deleting 8287 bp with paired sgRNAs^52^ in Nop2 gene N terminal (Figure S7D and S7E). Nop2 KD cells grew much slower in the presence of high MYC but similar to control in low MYC conditions (Figure S7F).

NOP2 has been proposed to regulate efficient ribosome biogenesis. We sought to determine if Nop2 was required for translation in EC4 cells. We detected nascent protein synthesis with the Click-iT HPG kit, and quantified GFP intensity in more than one thousand cells. Compared with MYC^high^ control cells, MYC^high^ *Nop2* KD cells had decreased protein synthesis (9105±4752 vs 5933±3530, p<0.000001). MYC^low^ *Nop2* KD cells had similar fluorescence intensity with MYC^low^ control cells (4178±1267 vs 4168±1433, p=0.832671) (Figure 5F and S7G).

By forming non-catalytic subcomplex with NIP7, NOP2 is required for ribosome assembly.^53^ To determine whether methyltransferase activity or non-catalytic scaffolding function of NOP2 was required for methionine dependence in MYC^high^ cells, it’s binding partner Nip7 was knocked down in EC4 cells by CRISPR (Figure S7H). Both MYC^high^ and MYC^low^ cell proliferation were significantly inhibited after *Nip7* knockdown (Figure 5G), indicating both MYC cell states were sensitive to the inhibition of ribosome assembly. Only MYC^high^ cells were potently inhibited after Nop2 depletion (Figure 5D, Figure S7F), this clearly demonstrated that the effect of Nop2 on MYC^high^ cells was because of methylation rather than non-catalytic ribosome assembly.

We also examined the DepMap project to check the dependency of liver cancer cell lines on NOP2. The dependence on a specific gene was measured by CERES, a computational method to estimate gene-dependency levels from CRISPR screens.^32^ The lower the score, the more likely a cell line was dependent on the gene. Liver cancer cell lines are dependent on methyltransferases FBL and NOP2, while DNMT1 was identified as a partially dependent gene (Figure S8A). NOP2 gene dependency highly correlated with MYC expression, meaning high MYC liver cancer cells were more dependent on NOP2 gene (Figure S8B). Other methyltransferases, including FBL, did not correlate with MYC gene levels. Not only MYC gene expression, many MYC up-regulated gene sets, including Hallmark MYC Targets v1 and v2, MYC oncogenic signature, MYC active pathway, were all negatively associated with NOP2 gene dependency, linking NOP2 depletion to diminished growth of cell lines with MYC-regulated gene sets in liver cancer cell lines (n=24) (Figure S8C).

### NOP2 depletion inhibits MYC-driven liver cancer

Given the selective growth inhibitory effects of NOP2 depletion in MYC high cells, we sought to determine if NOP2 depletion *in vivo* could limit liver tumor formation. Hydrodynamic delivery of nucleic acids into mouse liver allows for stable somatic engineering of hepatocytes.^54^ A mixture of 3 plasmids were injected, SB100 the transposase, pT3-MYC-IRES-Luc (transposon) to deliver MYC and luciferase, and pX330 to introduce Cas9 and a pair of *Nop2* sgRNAs, sgRNA 6 and 7 (Figure 6A). As a single sgRNA did not effectively deplete NOP2 protein expression, we used paired sgRNAs for the *in vivo* study. Two weeks after hydrodynamic injection, we visualized tumor growth via luciferase expression in the group with non-targeting sgRNA (Figure 6B); after 5-6 weeks, mice in the non-target (Ctrl) group reached the ethical endpoint and were sacrificed, and tissues were collected for analysis. Nop2 depletion markedly diminished MYC-driven tumorigenesis in the liver: all the mice in the Ctrl group developed large tumors; however, Nop2 gene-edited mice did not show evidence of macroscopic tumors (Figure 6C). Mice receiving the Nop2 sgRNA maintained a normal liver-to-body ratio (0.04) (Figure 6D). Tumor weight was not calculated as there were no discernible tumors after Nop2 gene editing. Immunohistochemistry staining showed both MYC and NOP2 highly expressed in tumors (Ctrl group), but their expressions were decreased in Nop2 gene editing group (Figure 6E). We also measured the survival of Nop2 CRISPR gene editing mice. Mice in the Ctrl group usually died of liver cancer between 5-7 weeks after hydrodynamic injection; mice in Nop2 gene editing group were monitored up to 12 weeks, twice the median survival time of the Ctrl group and all the mice survived this period and had histologically normal appearing livers (Figure 6F). After 3 months, Nop2 gene-edited mice still showed liver luciferase luminescence, suggesting their survival was due to Nop2 gene editing rather than loss of MYC-luciferase transgene expression (Figure 6G). These results demonstrated that knockdown of Nop2 selectively inhibited MYC high cell growth *in vitro* and prevented MYC liver tumor formation *in vivo*.

**Figure 6.**
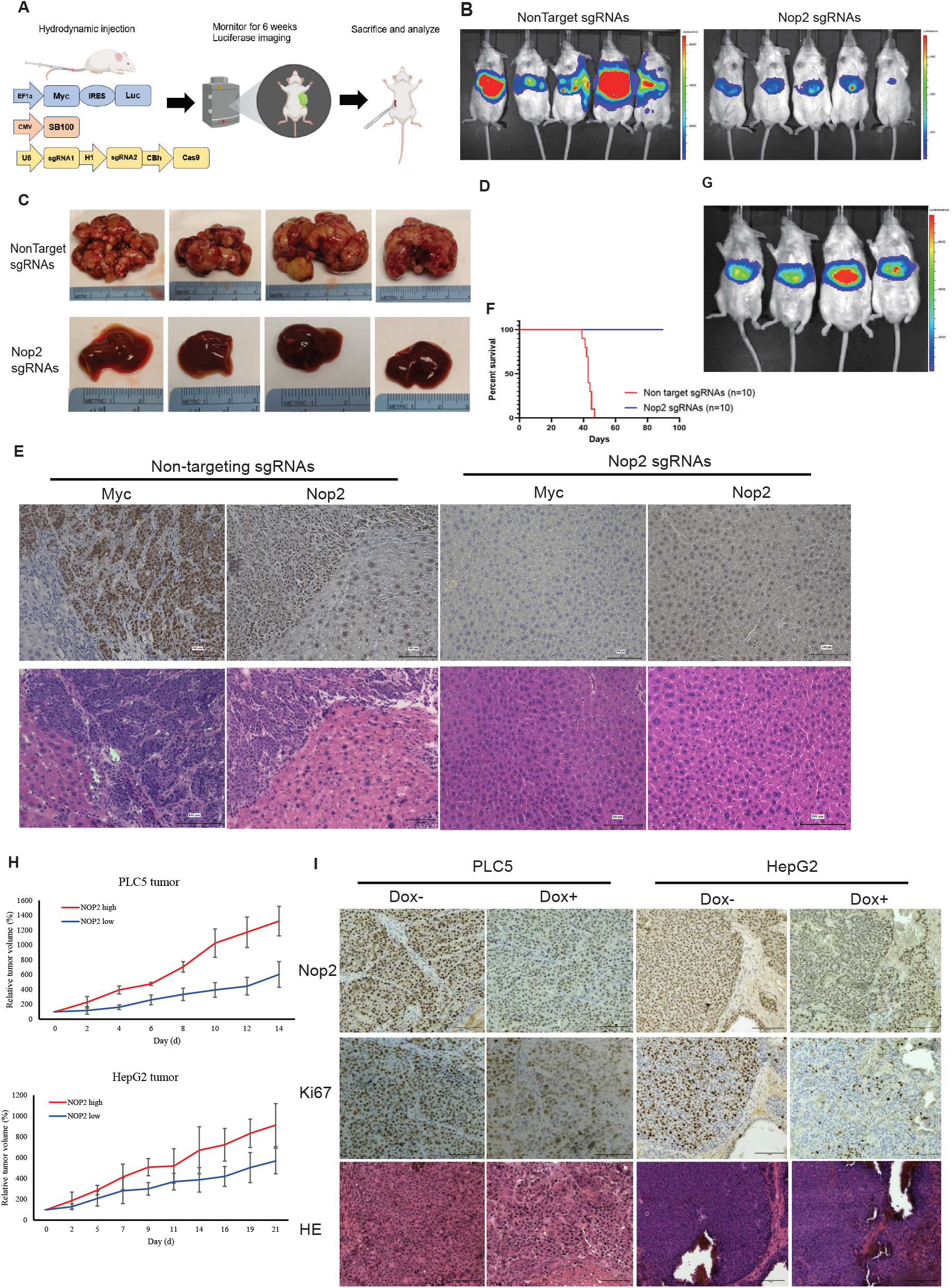
NOP2 depletion inhibits MYC-driven liver cancer *in vivo*. (A) Scheme of hydrodynamic injection of sleeping beauty transposon plasmids. *(B) in vivo* gene editing of Nop2. Two weeks after hydrodynamic injection, mice were imaged in IVIS. (C) Nop2 gene editing prevented MYC tumorigenesis *in vivo*. 6 weeks after hydrodynamic injection, mice were sacrificed and tumors/livers were shown. (D) Liver/body weight ratio in ctrl sgRNA group and Nop2 sgRNA group. (E) HE and IHC staining after Nop2 gene editing *in vivo*. Both Nop2 and Myc levels were decreased in Nop2 sgRNA group. Bar=100µm. (F) Survival of Nop2 gene editing mice and Ctrl mice (non-targeting sgRNAs). (G) Luciferase imaging of Nop2 gene editing mice 3 months after hydrodynamic injection. Nop2 gene editing mice still had luciferase fluorescence but no tumor was found. (H) Relative tumor volume of PLC5 and HepG2 xenografts. After tumor size reached 50-100 mm^3^, doxycycline food was used to induce the knockdown of NOP2. Tumor volume was measured every 2-3 days. (I) IHC staining of NOP2 and Ki67 in PLC5 and HepG2 tumors. Bar=100µm.

As a complementary approach, we delivered shRNA against mouse Nop2 *in vivo* to validate the effect of Nop2 knockdown on MYC-driven liver cancer (Figure S9A). Nop2 shRNA was made according to a commercialized Nop2 siRNA, Qiagen SI01328992, which was demonstrated to be effective in the body.^55^ Two plasmids were injected, pT3-MYC-U6-shNop2 and SB100, permitting co-expression of Nop2 shRNA with MYC. Similar to the CRISPR depletion studies, Nop2 shRNA also inhibited MYC-dependent tumor formation, and mice receiving shNop2 vector did not have macroscopic tumors (Figure S9B and S9C).

We next used an HCC xenograft cell line model to determine the dependence of NOP2 in MYC high HCC cell lines for tumor growth *in vivo*. HCC cells were infected with an inducible dCas9 vector pLX-TRE-dCas9 together with dual sgRNA plasmid expressing two sgRNAs targeting NOP2 promoter as reported.^56,57^ *In vitro* verification showed inducible CRISPRi knockdown worked in 4 of the 5 HCC cell lines (Figure S9D). Once subcutaneous tumor transplants reached 50-100 mm^3^, mice were fed doxycycline chow to induce NOP2 depletion. The ability of PLC/PRF/5 and HepG2 cells to form tumors in mouse models has been well-established, whereas Hep40 is much less tumorigenic. Both PLC5 and HepG2 tumors were significantly inhibited by CRISRRi knockdown of NOP2 (Figure 6H). Compared with HepG2, PLC5 cells had higher levels of NOP2 gene expression (Figure S5D) and MYC oncogenic signaling (Figure S5F); accordingly, the inhibition on PLC5 tumors was more effective. IHC staining confirmed decreased NOP2 protein expression in both tumors (Figure 6I). Likewise, the cell proliferation marker Ki67 staining was lower after NOP2 depletion, indicating diminished tumor growth proliferation. This further confirmed that NOP2 depletion in established tumors can substantially reduce human liver cancer growth.

## Discussion

Tumors use multiple metabolites to fuel their growth and cancer cells can reprogram metabolism due to activation of specific oncogenes. A favorable strategy to inhibit cancer is to restrict cancer cell access to specific nutrients critical for their growth. Beyond glucose and glutamine, serine, lipids, essential or non-essential amino acids (such as aspartic acid), all can be used as alternative sources to generate metabolic intermediates and compensate for the loss of a single nutrient source. Due to their metabolic plasticity, cancer cells may adjust their metabolic phenotypes to adapt and survive in different environments and conditions.^58,59^ Such metabolism plasticity makes the inhibition of just one nutrient source unable to curb cancer growth.

Carbon catabolism in cancer cells is much less redundant, with metabolic intermediates used for macromolecular synthesis to support cell growth, including nucleotide, cell membrane and protein synthesis. Methylation catabolizes carbon and relies predominantly on methionine metabolism, with SAM as the dominant methyl donor. Other methyl donors, such as folate, betaine, choline, and vitamin cofactors B2, B6, B12 and zinc, all rely on methionine metabolism. Thus, methionine metabolism is a metabolic bottleneck that rapidly growing cancer cells cannot readily bypass. Here we find that suppression of methionine metabolism through dietary, genetic or pharmacological strategies was efficient in blocking MYC-driven cell proliferation, *in vivo* tumor formation and growth. We identify Nop2, an rRNA methyltransferase, as a new target whose depletion selectively arrested MYC^high^ cell proliferation, prevented MYC-driven tumorigenesis and inhibited established cancer growth in mice.

Here we find that culturing MYC high liver cancer cells in low methionine reduced 28S rRNA methylation at three sites. In mouse cells NSUN5 is responsible for the methylation of the C3438 position,^60^ this may suggest that NSUN5 also plays a role in the methionine dependence of liver cancer cells. Both NOP2 and NSUN5 are methyltransferases of the NOL1/NOP2/SUN domain (NSUN) family, and share much similarity in catalytic domain.^61^ NSUN5 is reported to increase protein synthesis, cell proliferation and differentiation,^62^ and also regulates β-catenin mRNA degradation, leading to enhanced phagocytosis of tumor-associated macrophages and immune evasion.^63^ In HCC, NSUN5 plays an oncogenic role by stimulating ZBED3/Wnt/β-catenin signaling pathway.^64^ Interestingly, in MYC^low^ cells low methionine reduced C3438 methylation levels from ~80% to ~72%, while in MYC^high^ cells the decrease of C3438 methylation was much greater (from ~80% to ~55%), indicating MYC may contribute to the regulation on NSUN5 enzymic activity. We didn’t find that MYC induction led to NSUN5 transcriptional activation in the LT2-MYC model, so it may be an indirect effect. What is the function of NSUN5 in methionine dependence, and how MYC regulate NSUN5 methyltransferase activity, remain to be elucidated.

While cell culture experiments showed that depletion of NOP2 did not affect MYC protein expression (Figure 3H), here we observed Nop2 gene editing did prevent MYC driven liver cancer, thus reduced MYC protein levels *in vivo* (Figure 6E, Figure S9C). This discrepancy may reflect the fact that Nop2 depletion prevented tumorigenesis in naive liver. Likewise, MYC is known as a master regulator of ribosomal biogenesis, while ribosome genes are also feedback regulators of Myc.^65,66^ In the long-term, NOP2 deletion might diminished MYC expression via a feedback mechanism perhaps because cells are prevented from proliferating.

Finally, we found that NOP2 is important for efficient translation in MYC-high cells, whereas cells with lower MYC expression are less sensitive to NOP2 depletion, suggesting selective toxicity to MYC oncogene-overexpressing cells. Beyond rRNA processing, a recent paper^67^ demonstrated that NOP2 also regulates cell proliferation, rRNA processing and 60S ribosome biogenesis. Thus, there may be other mechanisms beyond m^5^C methylation of 28S rRNA that mediate some of the effects of NOP2 depletion in MYC high liver tumors. The authors in this recent study performed eCLIP-seq (GSE188735) to detect NOP2 binding to RNA fragments, in wild type and a C459A mutant (a mutant that stabilized NOP2 binding with RNA) in HEK293T cells. Surprisingly, although the majority of reads came from ribosome (rRNA, tRNA and snoRNAs), NOP2 was also found to bind to RNAs related to metabolism (AAMDC), signal transduction (SMAD9), DNA recombination (HFM1), splicing (RBM10), etc. Thus, NOP2 might have multiple cellular functions, providing a link between transcription and translation programs. While further research is needed to elucidate the function of NOP2 in cancer, our studies reveal that its inhibition may be a promising therapeutic target for MYC high liver tumors.

## Conflict of Interest

The authors have no competing interests.

## Author contributions

S.L. designed and performed the experiments, analyzed the data and wrote the manuscript. C.B. and D.K.N. designed and performed the metabolomics studies. M.S. performed the methionine tracing studies. X.L., X.C. and J.L. assisted with transgenic animal studies. V.R helped with computational analyses. A.G. supervised the project, designed the experiments. S.L. and A.G. wrote the manuscript.

## Acknowledgements

We thank Y. Zhou and I.H. Jain from Gladstone Institutes for data analysis of methionine tracing studies; M.H. Barcellos-Hoff and W. Chou from department of radiation oncology, UCSF and Helen Diller family comprehensive cancer center for assistance with *in vivo* imaging; A. Sil and A. Cohen from department of microbiology and immunology, UCSF to support with transgenic mouse models; J. Klefström from University of Helsinki for critical reading of the manuscript. The work was funded by NIH (R01CA223817 to A.G.), Marcus Program Transformative Integrated Research Award, and the Mark Foundation and Emerson Collective.

## Notes

### Competing Interest Statement

The authors have declared no competing interest.

