## Supplemental Data for "Methionine metabolism and the NOP2 methyltransferase are essential for MYC-Driven liver tumorigenesis"

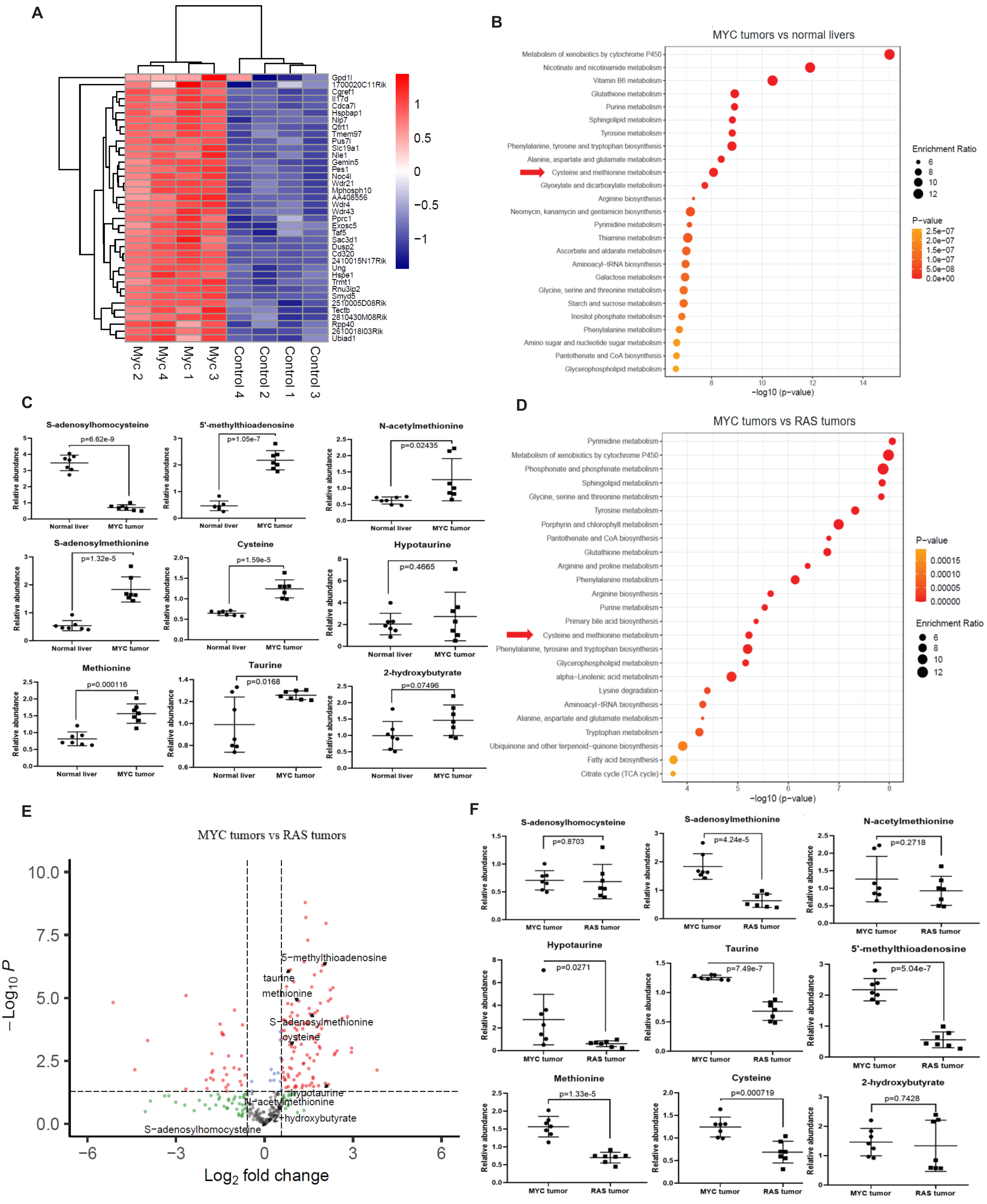
A
Gpd1l
1700020C11Rik
Cgref1
Il17d
Cdca7l
Hspbap1
Nip7
Qtrt1
Tmem97
Pus7l
Slc19a1
Nle1
Gemin5
Pes1
Noc4l
Wdr21
Mphosph10
AA408556
Wdr4
Wdr43
Pprc1
Exosc5
Taf5
Sac3d1
Dusp2
Cd320
2410015N17Rik
Ung
Hspe1
Trmt1
Rnu3ip2
Smyd5
2510005D08Rik
Tectb
2810430M08Rik
Rpp40
2610018I03Rik
Ubiad1
Myc 2
Myc 4
Myc 1
Myc 3
Control 4
Control 2
Control 1
Control 3
B
MYC tumors vs normal livers
Metabolism of xenobiotics by cytochrome P450
Nicotinate and nicotinamide metabolism
Vitamin B6 metabolism
Glutathione metabolism
Purine metabolism
Sphingolipid metabolism
Tyrosine metabolism
Phenylalanine, tyrosine and tryptophan biosynthesis
Alanine, aspartate and glutamate metabolism
Cysteine and methionine metabolism
Glyoxylate and dicarboxylate metabolism
Arginine biosynthesis
Neomycin, kanamycin and gentamicin biosynthesis
Pyrimidine metabolism
Thiamine metabolism
Ascorbate and aldarate metabolism
Aminoacyl−tRNA biosynthesis
Galactose metabolism
Glycine, serine and threonine metabolism
Starch and sucrose metabolism
Inositol phosphate metabolism
Phenylalanine metabolism
Amino sugar and nucleotide sugar metabolism
Pantothenate and CoA biosynthesis
Glycerophospholipid metabolism
−log10 (p−value)
Enrichment Ratio
P−value
C
S-adenosylhomocysteine
p=6.62e-9
5'-methylthioadenosine
p=1.05e-7
N-acetylmethionine
p=0.02435
S-adenosylmethionine
p=1.32e-5
Cysteine
p=1.59e-5
Hypotaurine
p=0.4665
Methionine
p=0.000116
Taurine
p=0.0168
2-hydroxybutyrate
p=0.07496
Relative abundance
Normal liver
MYC tumor
D
MYC tumors vs RAS tumors
Pyrimidine metabolism
Metabolism of xenobiotics by cytochrome P450
Phosphonate and phosphinate metabolism
Sphingolipid metabolism
Glycine, serine and threonine metabolism
Tyrosine metabolism
Porphyrin and chlorophyll metabolism
Pantothenate and CoA biosynthesis
Glutathione metabolism
Arginine and proline metabolism
Phenylalanine metabolism
Arginine biosynthesis
Purine metabolism
Primary bile acid biosynthesis
Cysteine and methionine metabolism
Phenylalanine, tyrosine and tryptophan biosynthesis
Glycerophospholipid metabolism
alpha−Linolenic acid metabolism
Lysine degradation
Aminoacyl−tRNA biosynthesis
Alanine, aspartate and glutamate metabolism
Tryptophan metabolism
Ubiquinone and other terpenoid−quinone biosynthesis
Fatty acid biosynthesis
Citrate cycle (TCA cycle)
−log10 (p−value)
P−value
Enrichment Ratio
E
MYC tumors vs RAS tumors
5−methylthioadenosine
taurine
methionine
S−adenosylmethionine
cysteine
hypotaurine
N−acetylmethionine
2−hydroxybutyrate
S−adenosylhomocysteine
− Log10 P
Log2 fold change
F
S-adenosylhomocysteine
p=0.8703
S-adenosylmethionine
p=4.24e-5
N-acetylmethionine
p=0.2718
Hypotaurine
p=0.0271
Taurine
p=7.49e-7
5'-methylthioadenosine
p=5.04e-7
Methionine
p=1.33e-5
Cysteine
p=0.000719
2-hydroxybutyrate
p=0.7428
Relative abundance
MYC tumor
RAS tumor

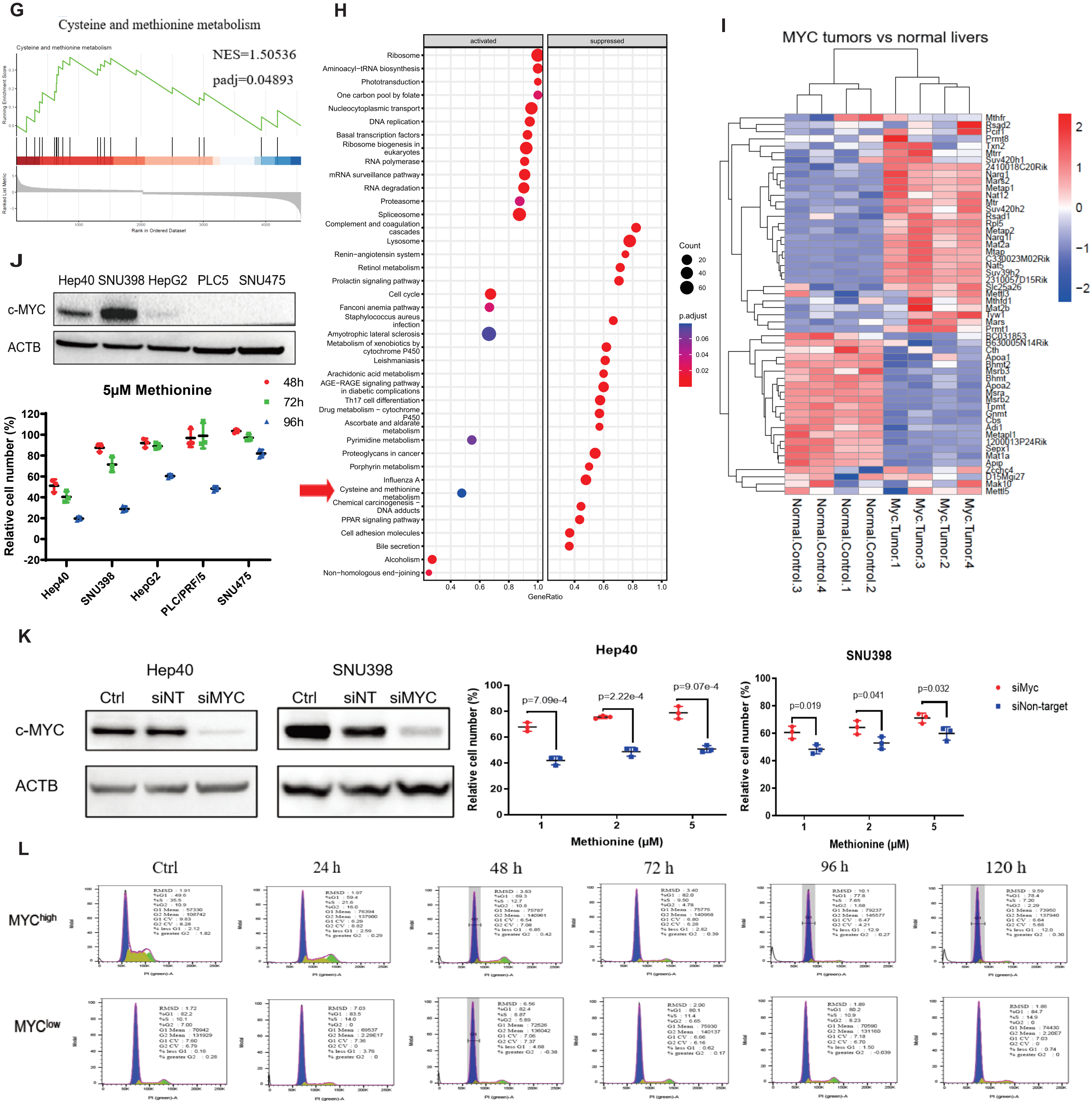
G
Cysteine and methionine metabolism
NES=1.50536
padj=0.04893
Running Enrichment Score
Ranked List Metric
Rank in Ordered Dataset
H
activated
suppressed
Ribosome
Aminoacyl−tRNA biosynthesis
Phototransduction
One carbon pool by folate
Nucleocytoplasmic transport
DNA replication
Basal transcription factors
Ribosome biogenesis in eukaryotes
RNA polymerase
mRNA surveillance pathway
RNA degradation
Proteasome
Spliceosome
Complement and coagulation cascades
Lysosome
Renin−angiotensin system
Retinol metabolism
Prolactin signaling pathway
Cell cycle
Fanconi anemia pathway
Staphylococcus aureus infection
Amyotrophic lateral sclerosis
Metabolism of xenobiotics by cytochrome P450
Leishmaniasis
Arachidonic acid metabolism
AGE−RAGE signaling pathway in diabetic complications
Th17 cell differentiation
Drug metabolism − cytochrome P450
Ascorbate and aldarate metabolism
Pyrimidine metabolism
Proteoglycans in cancer
Porphyrin metabolism
Influenza A
Cysteine and methionine metabolism
Chemical carcinogenesis − DNA adducts
PPAR signaling pathway
Cell adhesion molecules
Bile secretion
Alcoholism
Non−homologous end−joining
GeneRatio
Count
p.adjust
I
MYC tumors vs normal livers
Mthfr
Rsad2
Pcif1
Prmt8
Txn2
Mtrr
Suv420h1
2410018C20Rik
Narg1
Mars2
Metap1
Nat12
Mtr
Suv420h2
Rsad1
Rpl5
Metap2
Narg1l
Mat2a
Mtap
C330023M02Rik
Nat5
Suv39h2
2310057D15Rik
Slc25a26
Mettl3
Mthfd1
Mat2b
Tyw1
Mars
Prmt1
BC031853
B630005N14Rik
Cth
Apoa1
Bhmt2
Msrb3
Bhmt
Apoa2
Msra
Msrb2
Tpmt
Gnmt
Cbs
Adi1
Metapl1
1200013P24Rik
Sepx1
Mat1a
Apip
Zcchc4
D15Mgi27
Mak10
Mettl5
Normal.Control.3
Normal.Control.4
Normal.Control.1
Normal.Control.2
Myc.Tumor.1
Myc.Tumor.3
Myc.Tumor.2
Myc.Tumor.4
J
Hep40 SNU398 HepG2 PLC5 SNU475
c-MYC
ACTB
5μM Methionine
48h
72h
96h
Relative cell number (%)
Hep40
SNU398
HepG2
PLC/PRF/5
SNU475
K
Hep40
SNU398
Ctrl siNT siMYC
c-MYC
ACTB
Hep40
p=7.09e-4
p=2.22e-4
p=9.07e-4
SNU398
p=0.019
p=0.041
p=0.032
siMyc
siNon-target
Relative cell number (%)
Methionine (μM)
L
Ctrl
24 h
48 h
72 h
96 h
120 h
MYChigh
MYClow
PI (green)-A
Modal

**Figure S1. MYC-specific methionine metabolism alterations.**

(A) Heatmap of RNA seq showing relative expression genes known to be transactivated by MYC in LT2-MYC liver cancers versus control non-tumor liver.

(B) Pathway analysis showing enrichment of metabolites of MYC tumors as compared with normal livers. Arrow indicates cysteine and methionine metabolism.

(C) Methionine metabolism metabolite levels shown for MYC tumors vs normal livers.

(D) Pathway analysis showing enrichment of metabolites in MYC tumors as compared with RAS tumors. Arrow indicated cysteine and methionine metabolism.

(E) Volcano plot of metabolite alterations in MYC tumors vs RAS tumors. Methionine metabolism metabolites were shown.

(F) Methionine metabolism metabolite levels shown for MYC tumors vs RAS tumors.

(G) KEGG gene set cysteine and methionine metabolism was significantly enriched in $MYC^{high}$ cells ($MYC^{high}$ vs $MYC^{low}$).

(H) Dot plot of KEGG enrichment of gene expression in $MYC^{high}$ vs $MYC^{low}$ EC4 cells. Arrow indicates cysteine and methionine metabolism.

(I) Heatmap of methionine related genes in MYC tumors as compared with normal livers.

(J) Five human HCC cell lines were cultured with 5 μM methionine for 48-96 h, and cell proliferation was calculated by cell number counting. Cells cultured with full methionine medium (200μM) served as control.

(K) Knockdown of MYC attenuated relative growth inhibition induced by culture in low methionine for Hep40 and SNU398 cells. Cells were first treated with a pool of 10nM siRNAs against MYC for 48 h, then cultured with 1-5μM methionine medium or control in full media (200 μM) for another 72 h.

(L) Methionine starvation induced G1 cell cycle arrest only in EC4 $MYC^{high}$ but not $MYC^{low}$ cells.

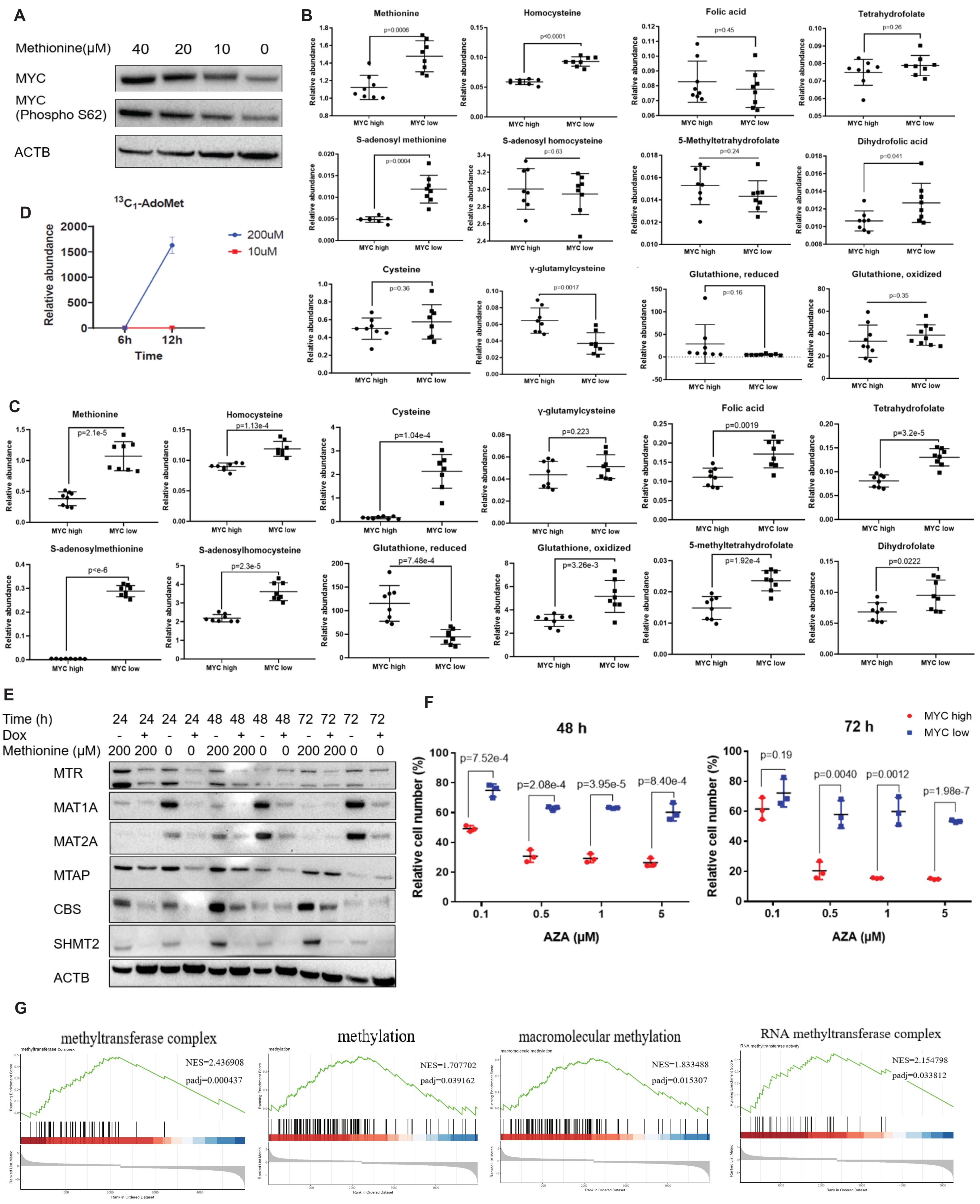
A
Methionine(μM) 40 20 10 0
MYC
MYC (Phospho S62)
ACTB
B
Methionine
p=0.0006
Homocysteine
p<0.0001
Folic acid
p=0.45
Tetrahydrofolate
p=0.26
S-adenosyl methionine
p=0.0004
S-adenosyl homocysteine
p=0.63
5-Methyltetrahydrofolate
p=0.24
Dihydrofolic acid
p=0.041
Cysteine
p=0.36
γ-glutamylcysteine
p=0.0017
Glutathione, reduced
p=0.16
Glutathione, oxidized
p=0.35
Relative abundance
MYC high
MYC low
D
13C1-AdoMet
200uM
10uM
Relative abundance
Time
6h
12h
C
Methionine
p=2.1e-5
Homocysteine
p=1.13e-4
Cysteine
p=1.04e-4
γ-glutamylcysteine
p=0.223
Folic acid
p=0.0019
Tetrahydrofolate
p=3.2e-5
S-adenosylmethionine
p<e-6
S-adenosylhomocysteine
p=2.3e-5
Glutathione, reduced
p=7.48e-4
Glutathione, oxidized
p=3.26e-3
5-methyltetrahydrofolate
p=1.92e-4
Dihydrofolate
p=0.0222
Relative abundance
MYC high
MYC low
E
Time (h) 24 24 24 24 48 48 48 48 72 72 72 72
Dox - + - + - + - + - + - +
Methionine (μM) 200 200 0 0 200 200 0 0 200 200 0 0
MTR
MAT1A
MAT2A
MTAP
CBS
SHMT2
ACTB
F
48 h
72 h
MYC high
MYC low
Relative cell number (%)
p=7.52e-4
p=2.08e-4
p=3.95e-5
p=8.40e-4
p=0.19
p=0.0040
p=0.0012
p=1.98e-7
AZA (μM)
0.1
0.5
1
5
G
methyltransferase complex
NES=2.436908
padj=0.000437
methylation
NES=1.707702
padj=0.039162
macromolecular methylation
NES=1.833488
padj=0.015307
RNA methyltransferase complex
NES=2.154798
padj=0.033812
Rank in Ordered Dataset

**Figure S2.** MYC-driven liver cancer is sensitive to methylation inhibition.

(A) Methionine regulated MYC expression and MYC S62 phosphorylation in EC4 cells.

(B) Levels of methionine metabolism metabolite in EC4 $MYC^{high}$ vs $MYC^{low}$ cells when cells were cultured with 20 µM methionine for 24 h (B) and 48 h (C).

(D) Methionine tracing study. Abundant 13C-SAM was detected in EC4 cells after cells were cultured with full methionine medium (200 µM) for 12h.

(E) Protein expression of key methionine metabolism enzymes in EC4 $MYC^{high}$ and $MYC^{low}$ cells maintained in either methionine free or methionine high (200µM) DMEM for 24-72 h. Data shown were representatives of at least 3 independent experiments.

(F) Dose-dependent effect of DNA/RNA methylation inhibitor AZA on EC4 cells. Inhibition was calculated by cell number counting.

(G) Enrichment of methyltransferase complex, methylation, macromolecular methylation and RNA methyltransferase complex in $MYC^{high}$ cells ($MYC^{high}$ vs $MYC^{low}$, RNA seq, enriched GO gene sets).

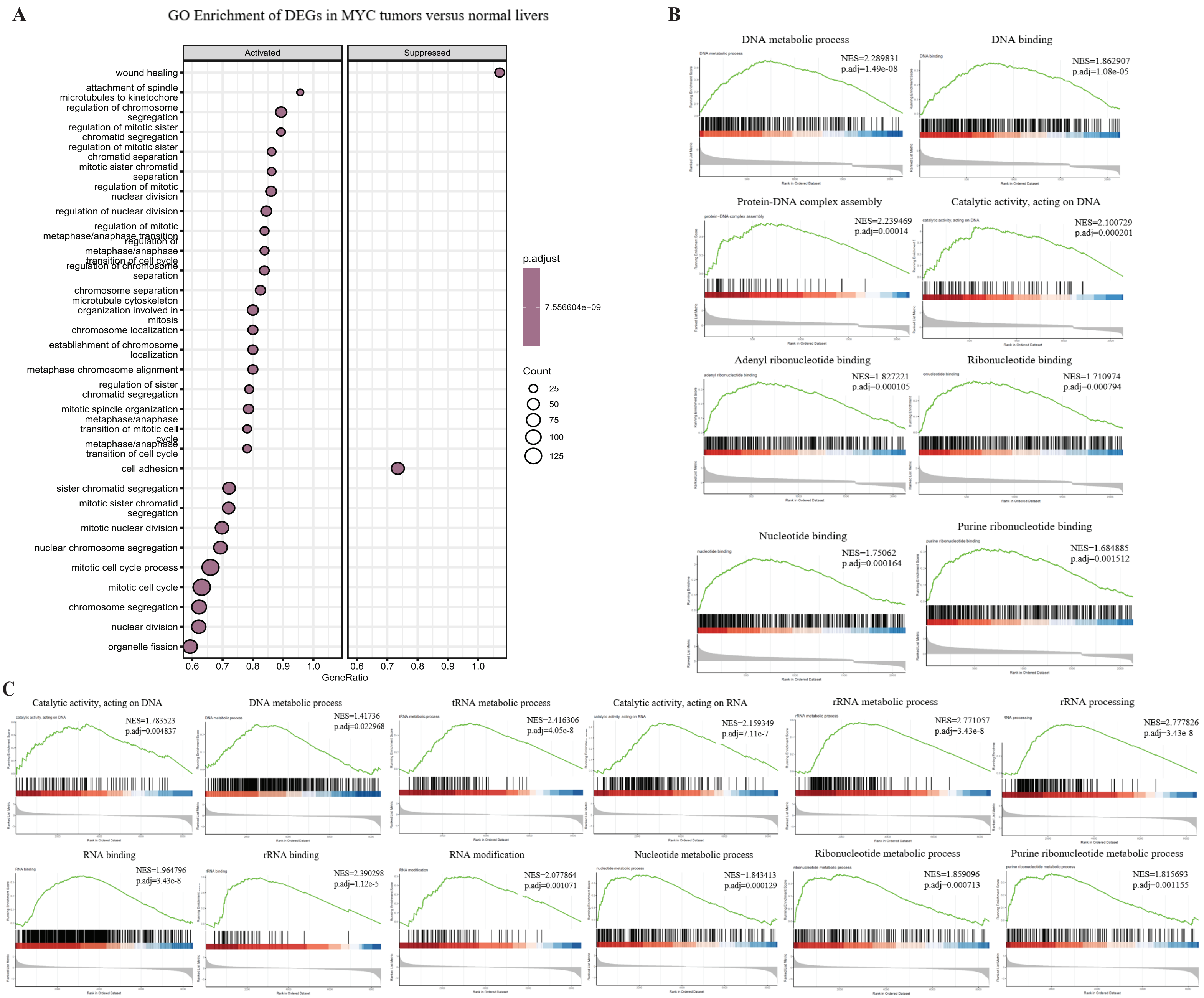
A
GO Enrichment of DEGs in MYC tumors versus normal livers
Activated
Suppressed
wound healing
attachment of spindle microtubules to kinetochore
regulation of chromosome segregation
regulation of mitotic sister chromatid segregation
regulation of mitotic sister chromatid separation
mitotic sister chromatid separation
regulation of mitotic nuclear division
regulation of nuclear division
regulation of mitotic metaphase/anaphase transition
regulation of metaphase/anaphase transition of cell cycle
regulation of chromosome separation
chromosome separation
microtubule cytoskeleton organization involved in mitosis
chromosome localization
establishment of chromosome localization
metaphase chromosome alignment
regulation of sister chromatid segregation
mitotic spindle organization
metaphase/anaphase transition of mitotic cell cycle
metaphase/anaphase transition of cell cycle
cell adhesion
sister chromatid segregation
mitotic sister chromatid segregation
mitotic nuclear division
nuclear chromosome segregation
mitotic cell cycle process
mitotic cell cycle
chromosome segregation
nuclear division
organelle fission
0.6 0.7 0.8 0.9 1.0
GeneRatio
p.adjust
7.556604e-09
Count
25
50
75
100
125
B
DNA metabolic process
NES=2.289831
p.adj=1.49e-08
DNA binding
NES=1.862907
p.adj=1.08e-05
Protein-DNA complex assembly
NES=2.239469
p.adj=0.00014
Catalytic activity, acting on DNA
NES=2.100729
p.adj=0.000201
Adenyl ribonucleotide binding
NES=1.827221
p.adj=0.000105
Ribonucleotide binding
NES=1.710974
p.adj=0.000794
Nucleotide binding
NES=1.75062
p.adj=0.000164
Purine ribonucleotide binding
NES=1.684885
p.adj=0.001512
C
Catalytic activity, acting on DNA
NES=1.783523
p.adj=0.004837
DNA metabolic process
NES=1.41736
p.adj=0.022968
tRNA metabolic process
NES=2.416306
p.adj=4.05e-8
Catalytic activity, acting on RNA
NES=2.159349
p.adj=7.11e-7
rRNA metabolic process
NES=2.771057
p.adj=3.43e-8
rRNA processing
NES=2.777826
p.adj=3.43e-8
RNA binding
NES=1.964796
p.adj=3.43e-8
rRNA binding
NES=2.390298
p.adj=1.12e-5
RNA modification
NES=2.077864
p.adj=0.001071
Nucleotide metabolic process
NES=1.843413
p.adj=0.000129
Ribonucleotide metabolic process
NES=1.859096
p.adj=0.000713
Purine ribonucleotide metabolic process
NES=1.815693
p.adj=0.001155

**Figure S3. Nucleotide modifications in MYC-driven liver cancer.**

(A) Go enrichment (dot plot) of DEGs in MYC tumors versus normal livers. Top 30 enriched gene sets were shown.

(B) Gene sets related to nucleotide modifications were significantly enriched in MYC tumors.

(C) Nucleotide modification gene sets were significantly enriched in $MYC^{high}$ cells ($MYC^{high}$ vs $MYC^{low}$, RNA seq, GO enrichment).

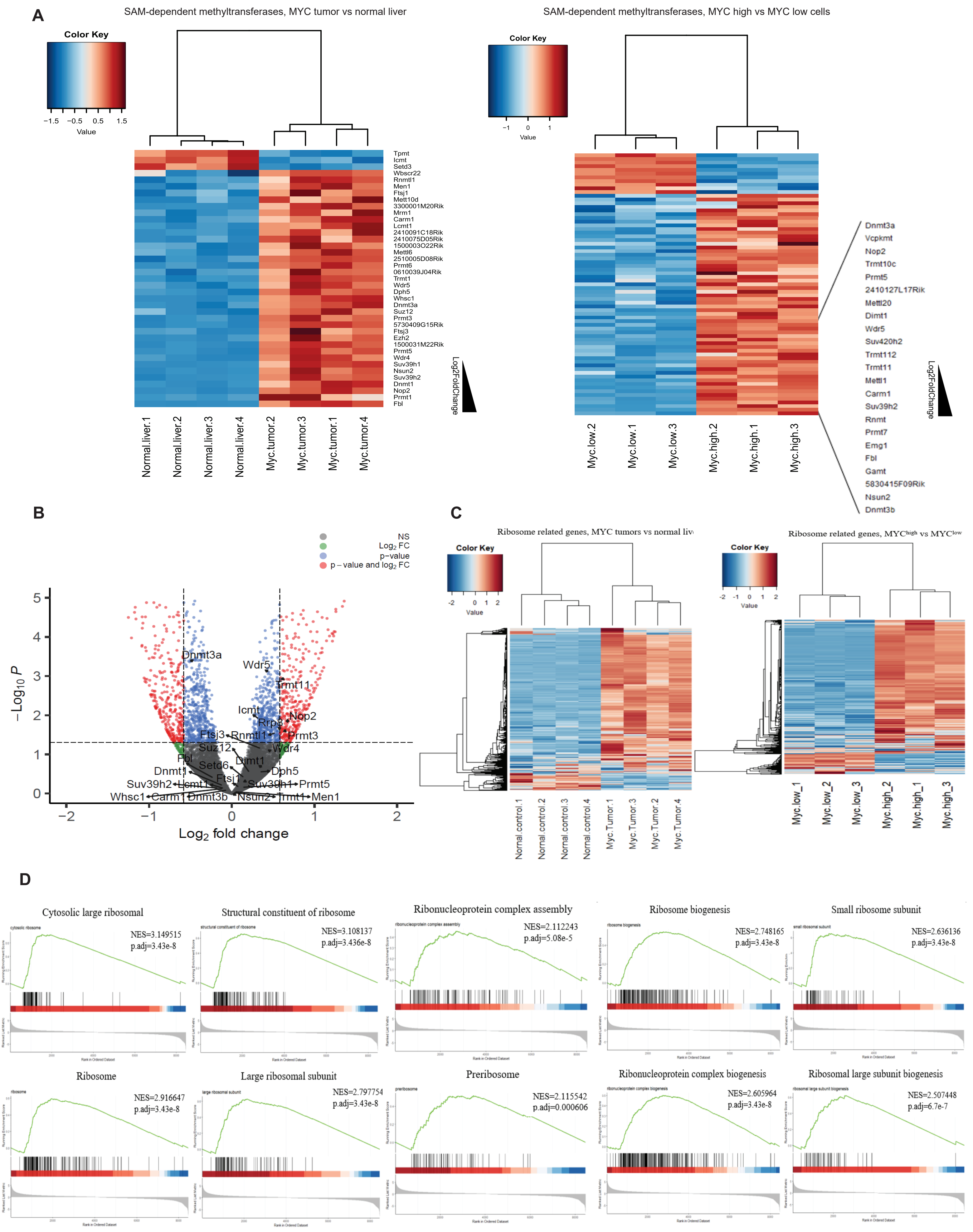
A
SAM-dependent methyltransferases, MYC tumor vs normal liver
SAM-dependent methyltransferases, MYC high vs MYC low cells
Color Key
Value
Tpmt
Icmt
Setd3
Wbscr22
Rnmtl1
Men1
Ftsj1
Mettl10d
3300001M20Rik
Mrm1
Carm1
Lcmt1
2410091C18Rik
2410075D05Rik
1500003O22Rik
Mettl6
2510005D08Rik
Prmt6
0610039J04Rik
Trmt1
Wdr5
Dph5
Whsc1
Dnmt3a
Suz12
Prmt3
5730409G15Rik
Ftsj3
Ezh2
1500031M22Rik
Prmt5
Wdr4
Suv39h1
Nsun2
Suv39h2
Dnmt1
Nop2
Prmt1
Fbl
Log2FoldChange
Normal.liver.1
Normal.liver.2
Normal.liver.3
Normal.liver.4
Myc.tumor.2
Myc.tumor.3
Myc.tumor.1
Myc.tumor.4
Dnmt3a
Vcpkmt
Nop2
Trmt10c
Prmt5
2410127L17Rik
Mettl20
Dimt1
Wdr5
Suv420h2
Trmt112
Trmt11
Mettl1
Carm1
Suv39h2
Rnmt
Prmt7
Emg1
Fbl
Gamt
5830415F09Rik
Nsun2
Dnmt3b
Myc.low.2
Myc.low.1
Myc.low.3
Myc.high.2
Myc.high.1
Myc.high.3
B
NS
Log2 FC
p−value
p − value and log2 FC
−Log10 P
Log2 fold change
Dnmt3a
Wdr5
Trmt11
Icmt
Nop2
Rrp9
Ftsj3
Rnmtl1
Prmt3
Suz12
Wdr4
Fbl
Setd6
Dimt1
Dnmt1
Dph5
Ftsj1
Suv39h2
Lcmt1
Suv39h1
Prmt5
Whsc1
Carm1
Dnmt3b
Nsun2
Trmt1
Men1
C
Ribosome related genes, MYC tumors vs normal liv
Ribosome related genes, MYChigh vs MYClow
Normal.control.1
Normal.control.2
Normal.control.3
Normal.control.4
Myc.Tumor.1
Myc.Tumor.3
Myc.Tumor.2
Myc.Tumor.4
Myc.low_1
Myc.low_2
Myc.low_3
Myc.high_2
Myc.high_1
Myc.high_3
D
Cytosolic large ribosomal
NES=3.149515
p.adj=3.43e-8
Structural constituent of ribosome
NES=3.108137
p.adj=3.436e-8
Ribonucleoprotein complex assembly
NES=2.112243
p.adj=5.08e-5
Ribosome biogenesis
NES=2.748165
p.adj=3.43e-8
Small ribosome subunit
NES=2.636136
p.adj=3.43e-8
Ribosome
NES=2.916647
p.adj=3.43e-8
Large ribosomal subunit
NES=2.797754
p.adj=3.43e-8
Preribosome
NES=2.115542
p.adj=0.000606
Ribonucleoprotein complex biogenesis
NES=2.605964
p.adj=3.43e-8
Ribosomal large subunit biogenesis
NES=2.507448
p.adj=6.7e-7

**Figure S4. Ribosome gene Nop2 was the most significantly altered m5C methyltransferase in LT2-MYC mouse model.**

(A) Heatmap of SAM-dependent methyltransferases in MYC tumors vs normal livers (microarray) and $MYC^{high}$ vs $MYC^{low}$ cells (RNA seq).

(B) Volcano plot of mRNA expression in MYC activated livers vs wild type livers (GSE83869). MYC was activated (dox off) for 48 h; after that mice were sacrificed and liver tissues were analyzed. The 26 methyltransferases upregulated both in MYC tumors and $MYC^{high}$ cells were shown.

(C) Heatmap of ribosome related genes in MYC tumors vs normal livers (left) and $MYC^{high}$ vs $MYC^{low}$ cells (right).

(D) Ribosome related gene sets were enriched in $MYC^{high}$ cells ($MYC^{high}$ vs $MYC^{low}$, RNA seq, GO enrichment).

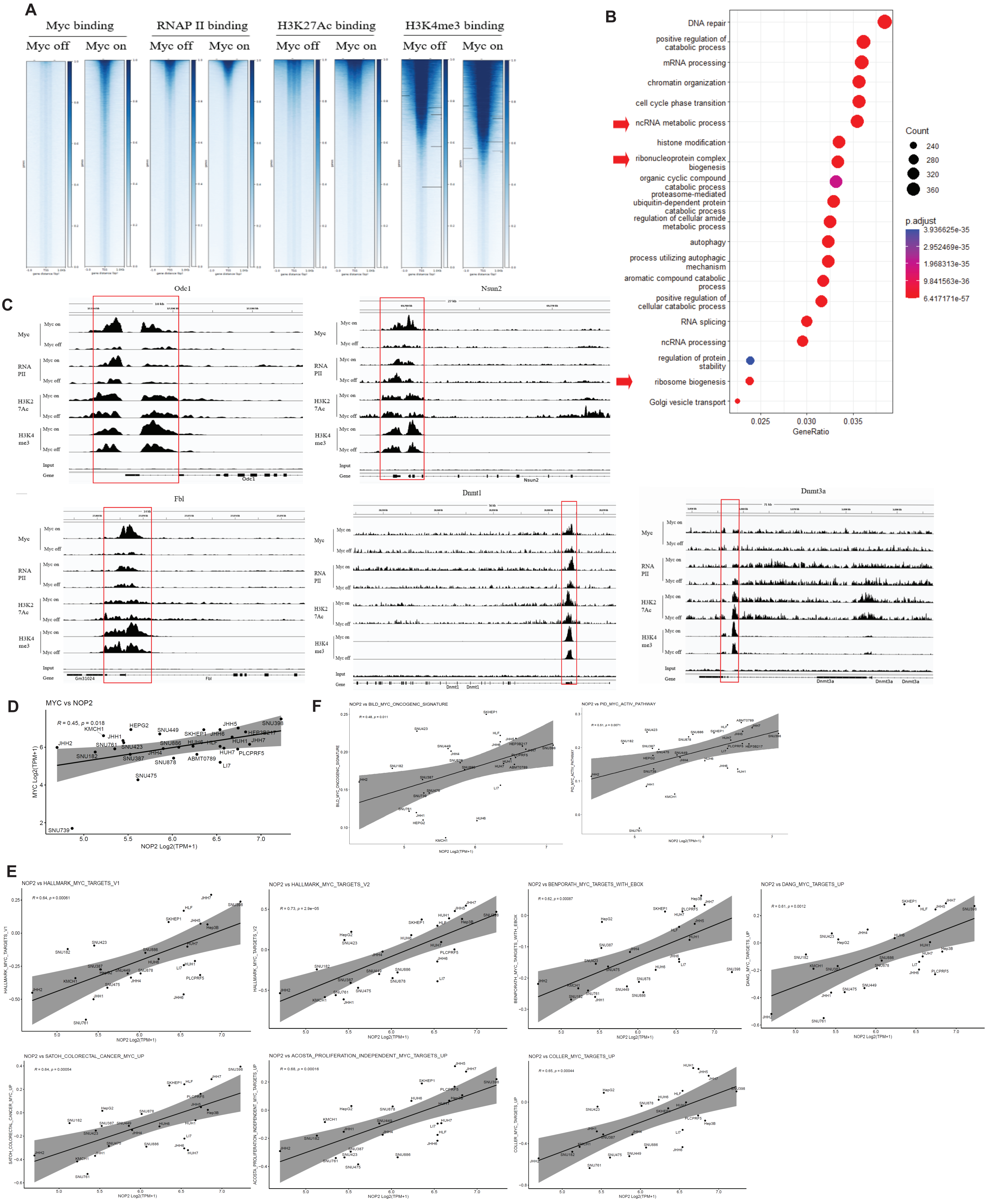
A
Myc binding
RNAP II binding
H3K27Ac binding
H3K4me3 binding
Myc off
Myc on
B
DNA repair
positive regulation of catabolic process
mRNA processing
chromatin organization
cell cycle phase transition
ncRNA metabolic process
histone modification
ribonucleoprotein complex biogenesis
organic cyclic compound catabolic process
proteasome-mediated ubiquitin-dependent protein catabolic process
regulation of cellular amide metabolic process
autophagy
process utilizing autophagic mechanism
aromatic compound catabolic process
positive regulation of cellular catabolic process
RNA splicing
ncRNA processing
regulation of protein stability
ribosome biogenesis
Golgi vesicle transport
0.025
0.030
0.035
GeneRatio
Count
240
280
320
360
p.adjust
3.936625e-35
2.952469e-35
1.968313e-35
9.841563e-36
6.417171e-57
C
Odc1
Nsun2
Fbl
Dnmt1
Dnmt3a
Myc
RNA PII
H3K27Ac
H3K4 me3
Myc on
Myc off
Input
Gene
D
MYC vs NOP2
R = 0.45, p = 0.018
MYC Log2(TPM+1)
NOP2 Log2(TPM+1)
F
NOP2 vs BILD_MYC_ONCOGENIC_SIGNATURE
R = 0.48, p = 0.011
NOP2 vs PID_MYC_ACTIV_PATHWAY
R = 0.51, p = 0.0071
E
NOP2 vs HALLMARK_MYC_TARGETS_V1
R = 0.64, p = 0.00061
NOP2 vs HALLMARK_MYC_TARGETS_V2
R = 0.73, p = 2.9e-05
NOP2 vs BENPORATH_MYC_TARGETS_WITH_EBOX
R = 0.62, p = 0.00087
NOP2 vs DANG_MYC_TARGETS_UP
R = 0.61, p = 0.0012
NOP2 vs SATOH_COLORECTAL_CANCER_MYC_UP
R = 0.64, p = 0.00054
NOP2 vs ACOSTA_PROLIFERATION_INDEPENDENT_MYC_TARGETS_UP
R = 0.68, p = 0.00016
NOP2 vs COLLER_MYC_TARGETS_UP
R = 0.65, p = 0.00044
NOP2 Log2(TPM+1)

**Figure S5. MYC binding on TSS sites in LT2-MYC transgenic mice liver.**

(A) Binding of Myc, RNAP II, H3K27Ac and H3K4me3 on mouse hepatocyte TSS sites (MYC ON versus control cells).

(B) Enrichment of MYC binding sites on TSS region (GSE83869).

(C) Specific binding of Myc, RNAP II, H3K27Ac and H3K4me3 on the promoter region of methyltransferases Fbl, Nsun2, Dnmt1 and Dnmt3. Odc1 served as positive control.

(D) The correlation between NOP2 and MYC gene expression (D), NOP2 and MYC target gene sets (E), NOP2 and MYC oncogenic signature (F) in 27 liver cancer cell lines (cancer dependency map project).

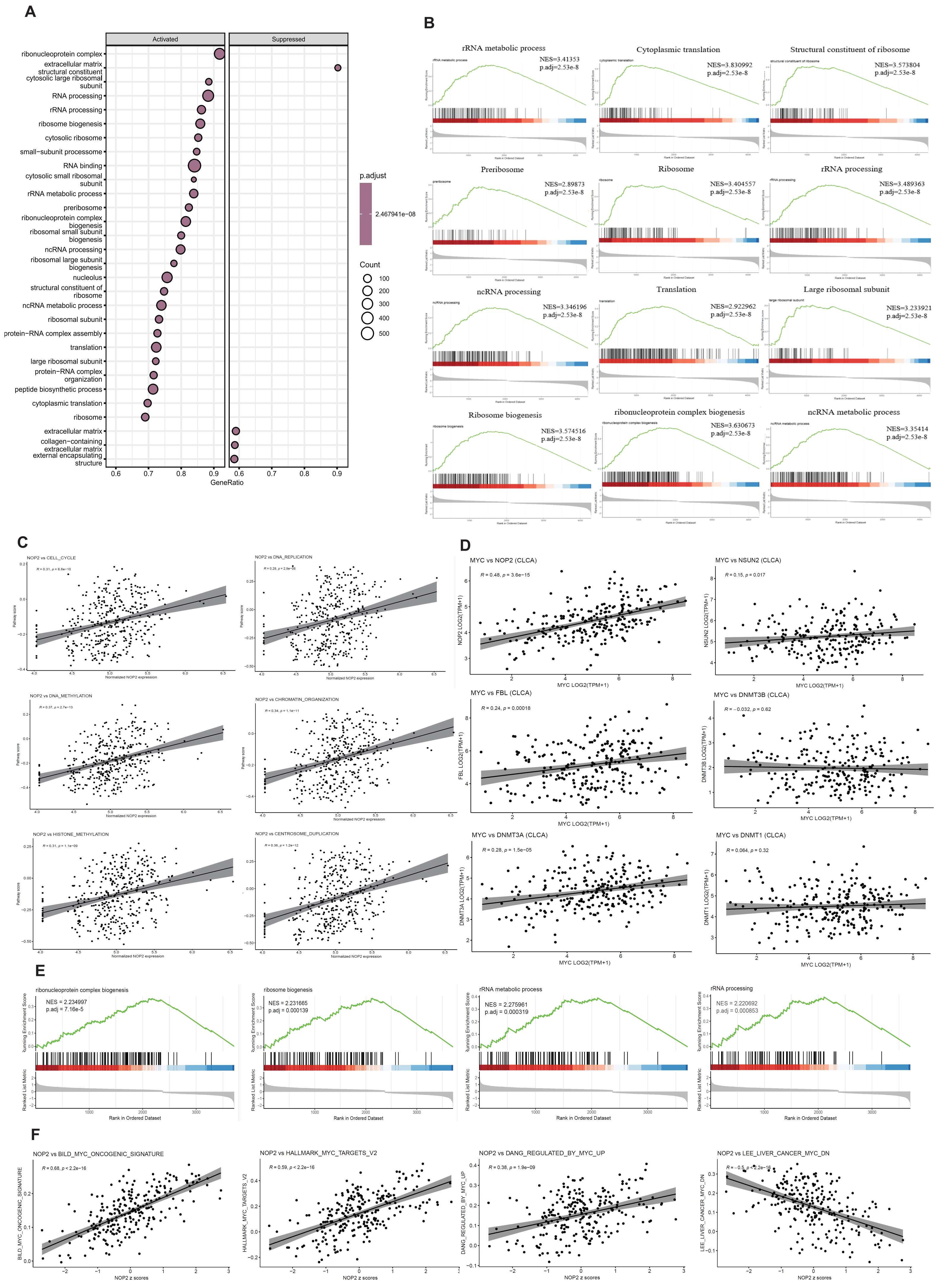
A
Activated
Suppressed
ribonucleoprotein complex
extracellular matrix structural constituent
cytosolic large ribosomal subunit
RNA processing
rRNA processing
ribosome biogenesis
cytosolic ribosome
small−subunit processome
RNA binding
cytosolic small ribosomal subunit
rRNA metabolic process
preribosome
ribonucleoprotein complex biogenesis
ribosomal small subunit biogenesis
ncRNA processing
ribosomal large subunit biogenesis
nucleolus
structural constituent of ribosome
ncRNA metabolic process
ribosomal subunit
protein−RNA complex assembly
translation
large ribosomal subunit
protein−RNA complex organization
peptide biosynthetic process
cytoplasmic translation
ribosome
extracellular matrix
collagen−containing extracellular matrix
external encapsulating structure
GeneRatio
p.adjust
2.467941e−08
Count
100
200
300
400
500
B
rRNA metabolic process NES=3.41353 p.adj=2.53e-8
Cytoplasmic translation NES=3.830992 p.adj=2.53e-8
Structural constituent of ribosome NES=3.573804 p.adj=2.53e-8
Preribosome NES=2.89873 p.adj=2.53e-8
Ribosome NES=3.404557 p.adj=2.53e-8
rRNA processing NES=3.489363 p.adj=2.53e-8
ncRNA processing NES=3.346196 p.adj=2.53e-8
Translation NES=2.922962 p.adj=2.53e-8
Large ribosomal subunit NES=3.233921 p.adj=2.53e-8
Ribosome biogenesis NES=3.574516 p.adj=2.53e-8
ribonucleoprotein complex biogenesis NES=3.630673 p.adj=2.53e-8
ncRNA metabolic process NES=3.35414 p.adj=2.53e-8
C
NOP2 vs CELL_CYCLE
NOP2 vs DNA_REPLICATION
NOP2 vs DNA_METHYLATION
NOP2 vs CHROMATIN_ORGANIZATION
NOP2 vs HISTONE_METHYLATION
NOP2 vs CENTROSOME_DUPLICATION
Normalized NOP2 expression
Pathway score
D
MYC vs NOP2 (CLCA) R = 0.48, p = 3.6e−15
MYC vs NSUN2 (CLCA) R = 0.15, p = 0.017
MYC vs FBL (CLCA) R = 0.24, p = 0.00018
MYC vs DNMT3B (CLCA) R = −0.032, p = 0.62
MYC vs DNMT3A (CLCA) R = 0.28, p = 1.5e−05
MYC vs DNMT1 (CLCA) R = 0.064, p = 0.32
MYC LOG2(TPM+1)
NOP2 LOG2(TPM+1)
NSUN2 LOG2(TPM+1)
FBL LOG2(TPM+1)
DNMT3B LOG2(TPM+1)
DNMT3A LOG2(TPM+1)
DNMT1 LOG2(TPM+1)
E
ribonucleoprotein complex biogenesis NES = 2.234997 p.adj = 7.16e-5
ribosome biogenesis NES = 2.231665 p.adj = 0.000139
rRNA metabolic process NES = 2.275961 p.adj = 0.000319
rRNA processing NES = 2.220692 p.adj = 0.000853
Running Enrichment Score
Ranked List Metric
Rank in Ordered Dataset
F
NOP2 vs BILD_MYC_ONCOGENIC_SIGNATURE R = 0.68, p < 2.2e−16
NOP2 vs HALLMARK_MYC_TARGETS_V2 R = 0.59, p < 2.2e−16
NOP2 vs DANG_REGULATED_BY_MYC_UP R = 0.38, p = 1.9e−09
NOP2 vs LEE_LIVER_CANCER_MYC_DN R = −0.5, p < 2.2e−16
BILD_MYC_ONCOGENIC_SIGNATURE
HALLMARK_MYC_TARGETS_V2
DANG_REGULATED_BY_MYC_UP
LEE_LIVER_CANCER_MYC_DN
NOP2 z scores

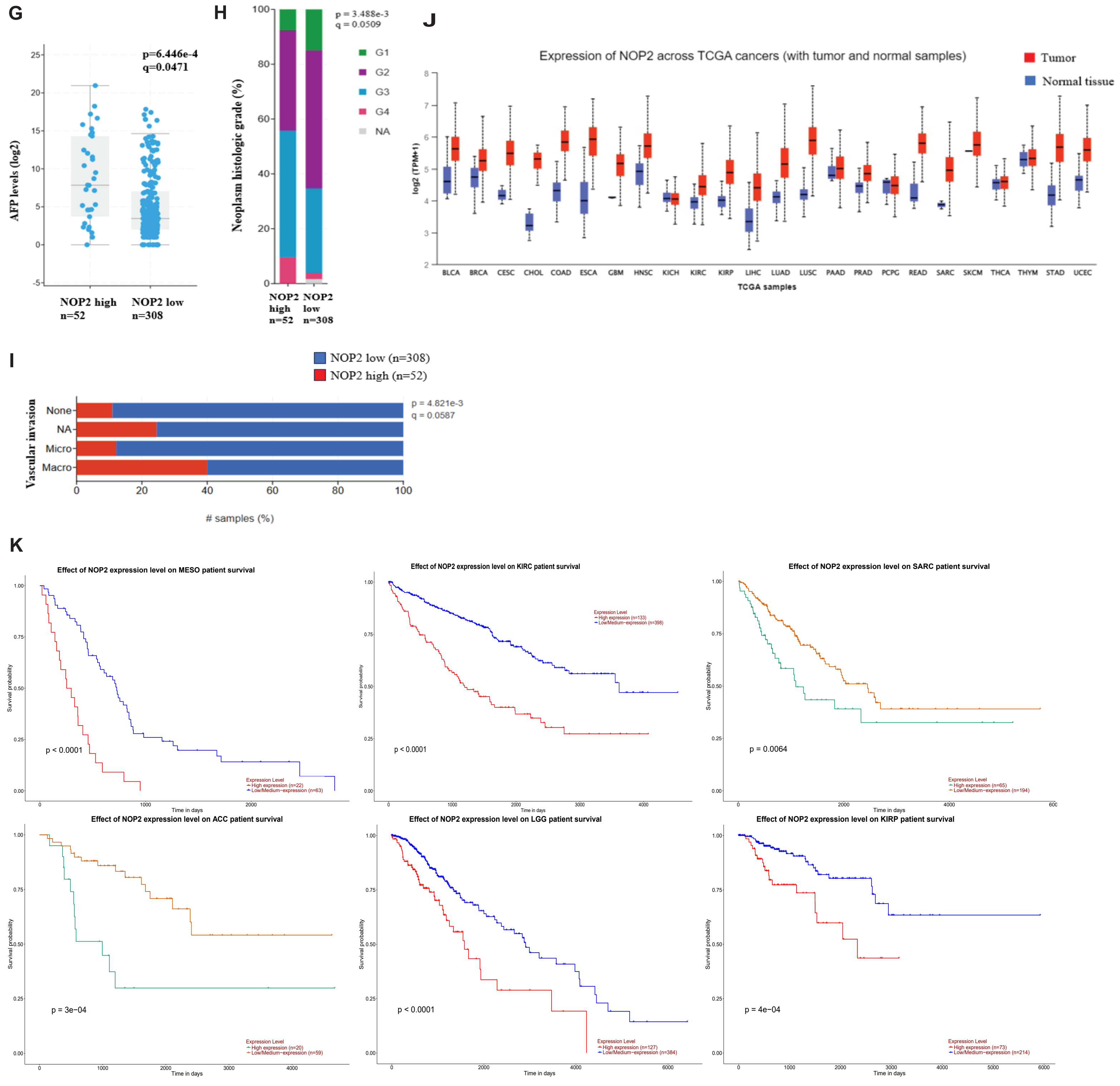
G
p=6.446e-4
q=0.0471
AFP levels (log2)
NOP2 high
n=52
NOP2 low
n=308
H
p = 3.488e-3
q = 0.0509
Neoplasm histologic grade (%)
G1
G2
G3
G4
NA
NOP2 high n=52
NOP2 low n=308
J
Expression of NOP2 across TCGA cancers (with tumor and normal samples)
Tumor
Normal tissue
log2 (TPM+1)
BLCA BRCA CESC CHOL COAD ESCA GBM HNSC KICH KIRC KIRP LIHC LUAD LUSC PAAD PRAD PCPG READ SARC SKCM THCA THYM STAD UCEC
TCGA samples
I
NOP2 low (n=308)
NOP2 high (n=52)
p = 4.821e-3
q = 0.0587
Vascular invasion
None
NA
Micro
Macro
### samples (%)
K
Effect of NOP2 expression level on MESO patient survival
p < 0.0001
Expression Level
High expression (n=22)
Low/Medium−expression (n=63)
Effect of NOP2 expression level on KIRC patient survival
Expression Level
High expression (n=133)
Low/Medium−expression (n=398)
p < 0.0001
Effect of NOP2 expression level on SARC patient survival
p = 0.0064
Expression Level
High expression (n=65)
Low/Medium−expression (n=194)
Effect of NOP2 expression level on ACC patient survival
p = 3e−04
Expression Level
High expression (n=20)
Low/Medium−expression (n=59)
Effect of NOP2 expression level on LGG patient survival
p < 0.0001
Expression Level
High expression (n=127)
Low/Medium−expression (n=384)
Effect of NOP2 expression level on KIRP patient survival
p = 4e−04
Expression Level
High expression (n=73)
Low/Medium−expression (n=214)
Survival probability
Time in days

**Figure S6. NOP2 associates with MYC and poor clinical outcome.**

(A) GO enrichment of TCGA liver cancer cohort, MYC high (n=61) versus MYC low (n=299).

(B) Ribosome related gene sets were top enriched in MYC high patients (TCGA cohort).

(C) ssGSEA analysis with TCGA liver cancer data, NOP2 gene expression associated with cell cycle, DNA replication, chromatin remodeling, histone and DNA methylation.

(D) The correlation between MYC and methyltransferases NOP2, FBL, NSUN2, DNMT1, DNMT3A, DNMT3B in CLCA cohort (n=239).

(E) Enrichment analysis in CLCA dataset. Ribosome biogenesis, rRNA processing, ribonucleoprotein complex biogenesis, rRNA processing were enriched in MYC high group.

(F) NOP2 gene expression highly associated with MYC signature and MYC target gene sets in CLCA cohort.

(G) AFP levels (G), neoplasm historical grade (H) and vascular invasion (I) were compared between NOP2 high group (mRNA expression > 85.56%, n=52) and NOP2 low group (mRNA expression ⩽ 85.56%, n=308).

(J) Pan-cancer expression of NOP2 in TCGA cohorts. NOP2 has higher levels in many cancers compared with normal tissue.

(K) Kaplan-Meier survival graph (NOP2 high vs NOP2 low) for mesothelioma (MESO), kidney renal clear cell carcinoma (KIRC), sarcoma (SARC), adrenocortical carcinoma (ACC), brain lower grade glioma (LGG) and kidney renal papillary cell carcinoma (KIRP) cancer patients.

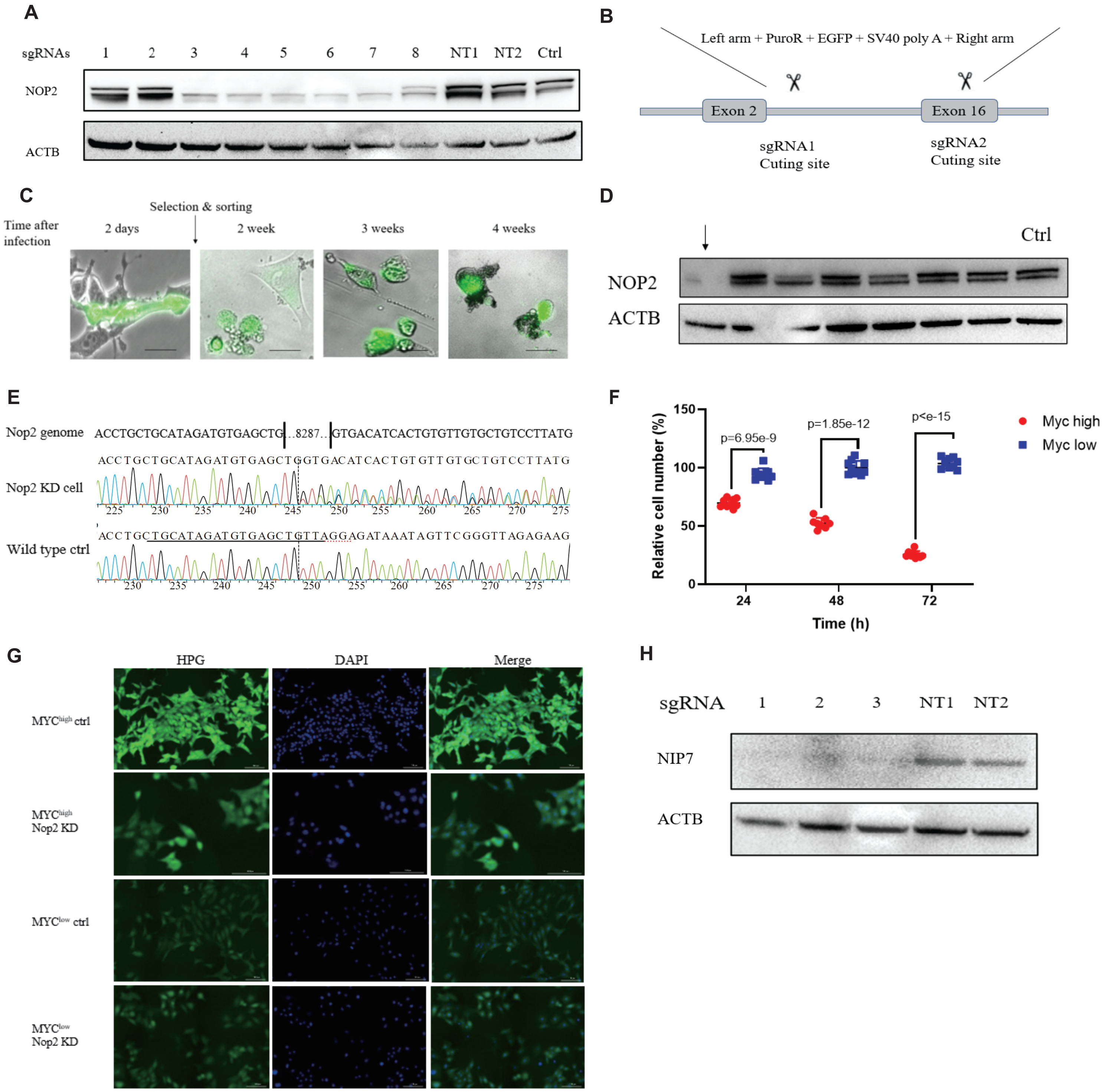
A
sgRNAs 1 2 3 4 5 6 7 8 NT1 NT2 Ctrl
NOP2
ACTB
B
Left arm + PuroR + EGFP + SV40 poly A + Right arm
Exon 2
Exon 16
sgRNA1
Cuting site
sgRNA2
Cuting site
C
Selection & sorting
Time after infection
2 days
2 week
3 weeks
4 weeks
D
Ctrl
NOP2
ACTB
E
Nop2 genome ACCTGCTGCATAGATGTGAGCTG…8287…GTGACATCACTGTGTTGTGCTGTCCTTATG
ACCTGCTGCATAGATGTGAGCTGGTGACATCACTGTGTTGTGCTGTCCTTATG
Nop2 KD cell
225 230 235 240 245 250 255 260 265 270 275
ACCTGCTGCATAGATGTGAGCTGTTAGGAGATAAATAGTTCGGGTTAGAGAAG
Wild type ctrl
230 235 240 245 250 255 260 265 270 275
F
Relative cell number (%)
150
100
50
0
p=6.95e-9
p=1.85e-12
p<e-15
Myc high
Myc low
24
48
72
Time (h)
G
HPG
DAPI
Merge
MYChigh ctrl
MYChigh Nop2 KD
MYClow ctrl
MYClow Nop2 KD
H
sgRNA 1 2 3 NT1 NT2
NIP7
ACTB

**Figure S7. Effect of Nop2 on MYC$^{high}$ and MYC$^{low}$ EC4 cells.**

(A) CRISPR knockdown in EC4 cells. Eells were transfected with different Nop2 sgRNA, further selected with puromycin for 3 days and then subjected to western blot.

(B) A scheme of pair sgRNAs to delete part of Nop2 gene in EC4 cells.

(C) MYC$^{high}$ cells cannot survive after complete Nop2 loss. Cells were infected with guide RNA virus harboring paired Nop2 sgRNAs, after 48h cells were selected and single-cell sorted into 96 well plates. Cells were maintained until Nop2 KO or KD clone appeared. Bar=10μm.

(D) Western blot confirmed Nop2 knockdown. Arrow pointed at the Nop2 KD cell line.

(E) Sequencing of the Nop2 KD cell line. More than 8000bp was deleted with Nop2 paired sgRNAs.

(F) Knockdown of Nop2 arrested MYC$^{high}$ cell growth (compared with normal MYC$^{high}$ cells). No difference between MYC$^{low}$ Nop2 KD cells and Ctrl MYC$^{low}$ cells were observed.

(G) Fluorescence imaging of MYC$^{high}$ Ctrl vs MYC$^{high}$ Nop2 KD cells, and MYC$^{low}$ Ctrl vs MYC$^{low}$ Nop2 KD cells. The Click-iT™ HPG protein synthesis assay was employed to measure nascent protein production. Bar=25μm.

(H) Effect of Nip7 sgRNAs on protein expression was measured by western blot.

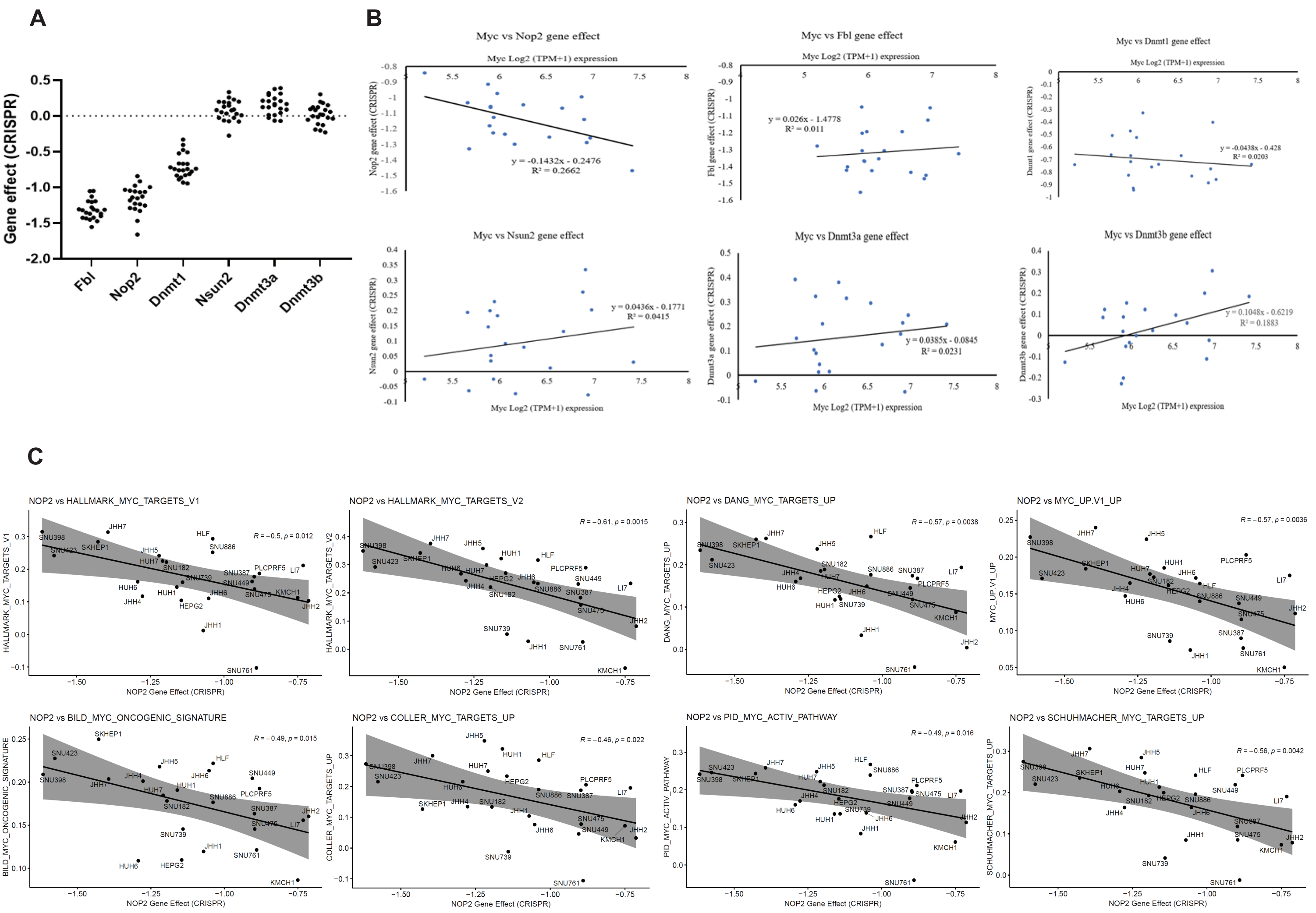
A
B
C
Gene effect (CRISPR)
Fbl
Nop2
Dnmt1
Nsun2
Dnmt3a
Dnmt3b
Myc vs Nop2 gene effect
y = -0.1432x - 0.2476
R² = 0.2662
Myc vs Fbl gene effect
y = 0.026x - 1.4778
R² = 0.011
Myc vs Dnmt1 gene effect
y = -0.0438x - 0.428
R² = 0.0203
Myc vs Nsun2 gene effect
y = 0.0436x - 0.1771
R² = 0.0415
Myc vs Dnmt3a gene effect
y = 0.0385x - 0.0845
R² = 0.0231
Myc vs Dnmt3b gene effect
y = 0.1048x - 0.6219
R² = 0.1883
Myc Log2 (TPM+1) expression
NOP2 vs HALLMARK_MYC_TARGETS_V1
R = −0.5, p = 0.012
NOP2 vs HALLMARK_MYC_TARGETS_V2
R = −0.61, p = 0.0015
NOP2 vs DANG_MYC_TARGETS_UP
R = −0.57, p = 0.0038
NOP2 vs MYC_UP.V1_UP
R = −0.57, p = 0.0036
NOP2 vs BILD_MYC_ONCOGENIC_SIGNATURE
R = −0.49, p = 0.015
NOP2 vs COLLER_MYC_TARGETS_UP
R = −0.46, p = 0.022
NOP2 vs PID_MYC_ACTIV_PATHWAY
R = −0.49, p = 0.016
NOP2 vs SCHUHMACHER_MYC_TARGETS_UP
R = −0.56, p = 0.0042
NOP2 Gene Effect (CRISPR)

**Figure S8. NOP2 dependency correlates with MYC and MYC target gene sets.**

(A) gene effect of m5C methyltransferases in liver cancer cell lines, CRISPR data sourced from CERES. Lower score means cell lines are more likely to dependent on a given gene. A score of 0 is equivalent to a gene that is not essential whereas a score of -1 corresponds to the median of all common essential genes.

(B) NOP2 gene effect negatively correlated with MYC gene expression in liver cancer cell lines, indicating MYC high liver cancer cell lines were more dependent on NOP2 than MYC low cells.

(C) NOP2 gene dependency negatively correlated with various MYC target gene sets. This suggested that liver cancer cell lines expressing high MYC target genes were more dependent on NOP2.

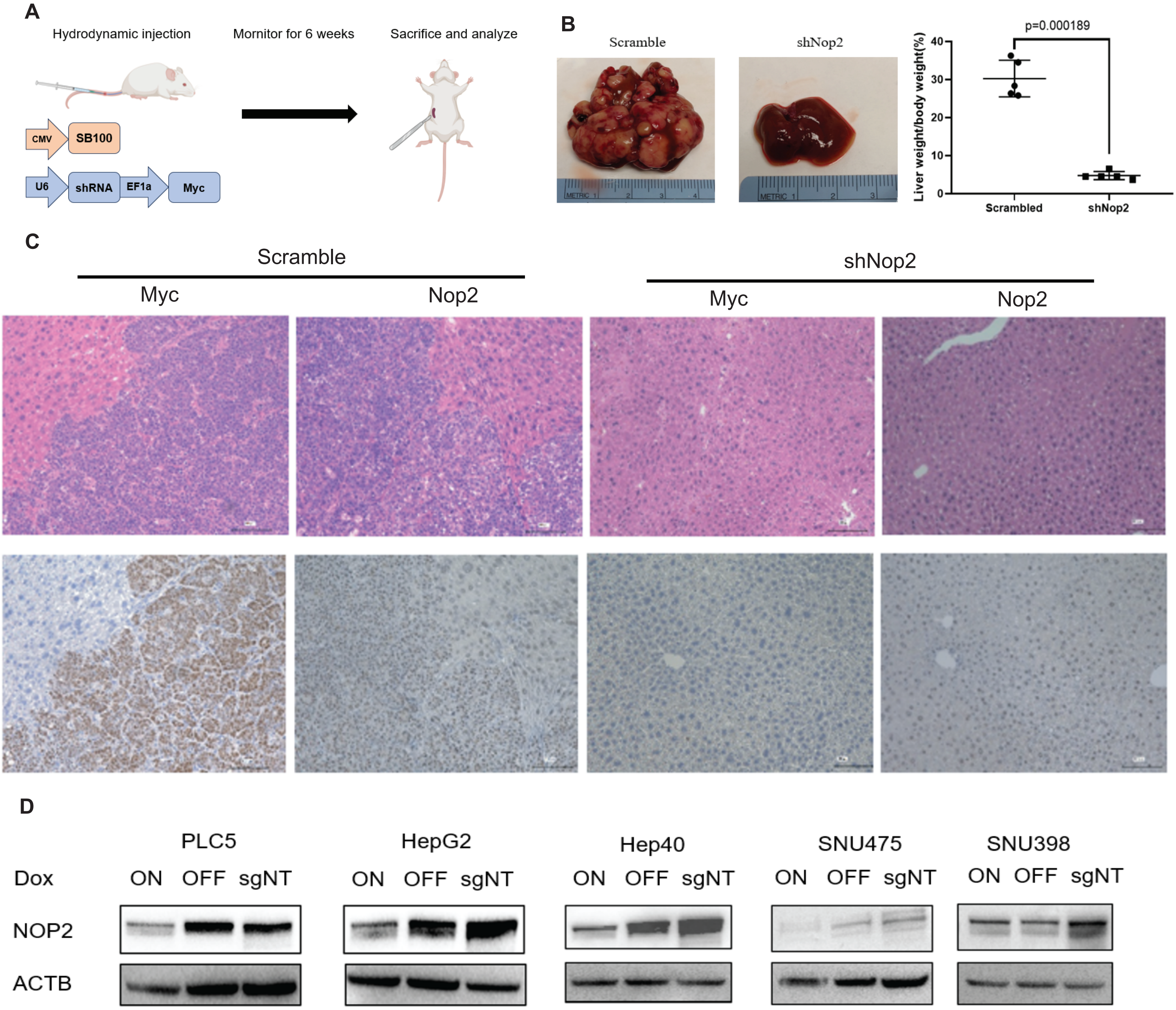
A
Hydrodynamic injection
Mornitor for 6 weeks
Sacrifice and analyze
CMV
SB100
U6
shRNA
EF1a
Myc
B
Scramble
shNop2
p=0.000189
Liver weight/body weight(%)
40
30
20
10
0
Scrambled
shNop2
C
Scramble
shNop2
Myc
Nop2
Myc
Nop2
D
PLC5
HepG2
Hep40
SNU475
SNU398
Dox
ON OFF sgNT
NOP2
ACTB

**Figure S9. NOP2 depletion inhibited MYC-driven liver cancer**

(A) Scheme of shNop2 animal study and sleeping beauty vectors.

(B) Representative mouse liver 6 weeks after hydrodynamic injection.

(C) H&E (top) and IHC staining (bottom) for Myc and Nop2 of liver samples. Scramble shRNA (ctrl) vs shNop2 are indicated. Bar=100μm.

(D) Inducible CRISPRi knockdown of NOP2 in HCC cell lines. Cells were transfected with pLX-TRE-dCas9 and pDECKO containing two sgRNAs targeting NOP2 promoter region, then treated with 1μg/ml doxycycline for 3-4 days. NOP2 protein was measured by western blot.
